# Broad ultra-potent neutralization of SARS-CoV-2 variants by monoclonal antibodies specific to the tip of RBD

**DOI:** 10.1101/2021.09.24.461616

**Authors:** Hang Ma, Yingying Guo, Haoneng Tang, Chien-Te K. Tseng, Lei Wang, Huifang Zong, Zhenyu Wang, Yang He, Yunsong Chang, Shusheng Wang, Haiqiu Huang, Yong Ke, Yunsheng Yuan, Mingyuan Wu, Yuanyuan Zhang, Aleksandra Drelich, Kempaiah Rayavara Kempaiah, Bi-Hung Peng, Ailin Wang, Kaiyong Yang, Haiyang Yin, Junjun Liu, Yali Yue, Wenbo Xu, Shuangli Zhu, Tianjiao Ji, Xiaoju Zhang, Ziqi Wang, Gang Li, Guangchun Liu, Jingjing Song, Lingling Mu, ZongShang Xiang, Zhangyi Song, Hua Chen, Yanlin Bian, Baohong Zhang, Hui Chen, Jiawei Zhang, Yunji Liao, Li Zhang, Li Yang, Yi Chen, John Gilly, Xiaodong Xiao, Lei Han, Hua Jiang, Yueqing Xie, Qiang Zhou, Jianwei Zhu

## Abstract

Severe acute respiratory syndrome coronavirus 2 (SARS-CoV-2) variants of concern (VOCs) continue to wreak havoc across the globe. Higher transmissibility and immunologic resistance of VOCs bring unprecedented challenges to epidemic extinguishment. Here we describe a monoclonal antibody, 2G1, that neutralizes all current VOCs and has surprising tolerance to mutations adjacent to or within its interaction epitope. Cryo-electron microscopy structure showed that 2G1 bound to the tip of receptor binding domain (RBD) of spike protein with small contact interface but strong hydrophobic effect, which resulted in nanomolar to sub-nanomolar affinities to spike proteins. The epitope of 2G1 on RBD partially overlaps with ACE2 interface, which gives 2G1 ability to block interaction between RBD and ACE2. The narrow binding epitope but high affinity bestow outstanding therapeutic efficacy upon 2G1 that neutralized VOCs with sub-nanomolar IC_50_ *in vitro*. In SARS-CoV-2 and Beta- and Delta-variant-challenged transgenic mice and rhesus macaque models, 2G1 protected animals from clinical illness and eliminated viral burden, without serious impact to animal safety. Mutagenesis experiments suggest that 2G1 could be potentially capable of dealing with emerging SARS-CoV-2 variants in future. This report characterized the therapeutic antibodies specific to the tip of spike against SARS-CoV-2 variants and highlights the potential clinical applications as well as for developing vaccine and cocktail therapy.

## Introduction

Since the first Coronavirus Disease 2019 (COVID-19) case was diagnosed at the end of 2019, the severe acute respiratory syndrome coronavirus 2 (SARS-CoV-2) has caused more than 200 million confirmed infections and 4.5 million deaths in the following eighteen months, with no sign of stopping (https://ourworldindata.org/coronavirus)^1–6^. The hope-placed distribution of vaccines once appeared effectively controlling the virus spread. However, the antigenic evolution of SARS-CoV-2, especially in the spike (S) protein associated with receptor binding, alters the viral immunogenicity facilitating the virus’s immune escape and crossing transmission barriers^7,8^.

Receptor binding domain (RBD) on the S protein is a determinant that mediates the binding of SARS-CoV-2 to the angiotensin-converting enzyme 2 (ACE2). Neutralizing antibodies targeting RBD were proved to be effective^9–11^. Correspondingly, substitutions on RBD may reduce neutralizing efficacy^12–14^. Several variants, listed as Variant of Concern (VOC), featured with RBD substitutions and non-RBD mutations showed to have higher transmissibility and led to more severe illness^15–17^, which has been causing great global dissemination concern. SARS-CoV-2 B.1.1.7 (Alpha) was first identified in United Kingdom in late summer of 2020 and rapidly became the dominant variant. This variant has nine mutations in the S protein, one of which is N501Y in RBD^18^. Alpha variant possesses a comparative transmission advantage, with a reproductive number 50% to 100% higher than other non-VOC lineages^1^. Vaccine-elicited neutralizing antibody responses were shown to be at risk of being desensitized by Alpha^19^. SARS-CoV-2 B.1.351 (Beta) has three substitutions in RBD, i.e., K417N, E484K, and N501Y. Incorporation of E484K empowers variants possible being completely resistant to plasma neutralization^20^. Mutations E484K together with K417N and N501Y largely contribute to the escape of Beta variant from convalescent and vaccine-induced sera^21,22^. SARS-CoV-2 P.1 (Gamma) shares three identical site-mutations in RBD with Beta variant, and their differences are that the substitution of K417 is threonine in Gamma variant, while is asparagine in Beta variant. Similarly, Gamma variant notably reduced susceptibility to antibody treatment and vaccine protection^23,24^. SARS-CoV-2 B.1.617.2 (Delta) was first reported in India and quickly spread globally in the first half of 2021. This strain has more than ten S protein mutations and two of them, L452R and T478K, are in RBD. Delta variant exhibits more extensive immunologic resistance than Alpha, escaping from many S protein antibodies targeting RBD and non-RBD epitopes^25,26^. Individuals who recovered from Beta and Gamma variants are more susceptible to be infected with Delta^27^. In addition to these VOCs, potential outbreaks of several variants have raised public concern, such as the recently rapidly spreading variant C.37 (Lambda)^28^ and the new variant B.1.621 (Mu)^29^. The emergence of these variants, even possible hybrid variants, raises the risk of compromising the therapeutic effectiveness of vaccines and neutralizing antibodies that were previously developed^30,31^.

Here we report our efforts on discovering neutralizing antibodies that provide extensive protection against the variants with global impact, especially the VOCs. We isolated RBD positive single B cells from convalescent individuals and cloned monoclonal antibodies (mAbs) within. After a series of programmed screening, several antibodies with remarkable neutralizing effect were panned out from the candidates (Fig. 1a). One of these antibodies, designated as 2G1, efficiently neutralized all VOCs including widely spread Alpha, Beta, Gamma, Delta variants and Cluster 5, a variant with Y453F substitution once caused public concern due to the zoonotic characteristics. The antibody 2G1 was subsequently fully characterized physic-chemically and biologically, as well as evaluated in potential in clinical applications.

**Fig. 1.**
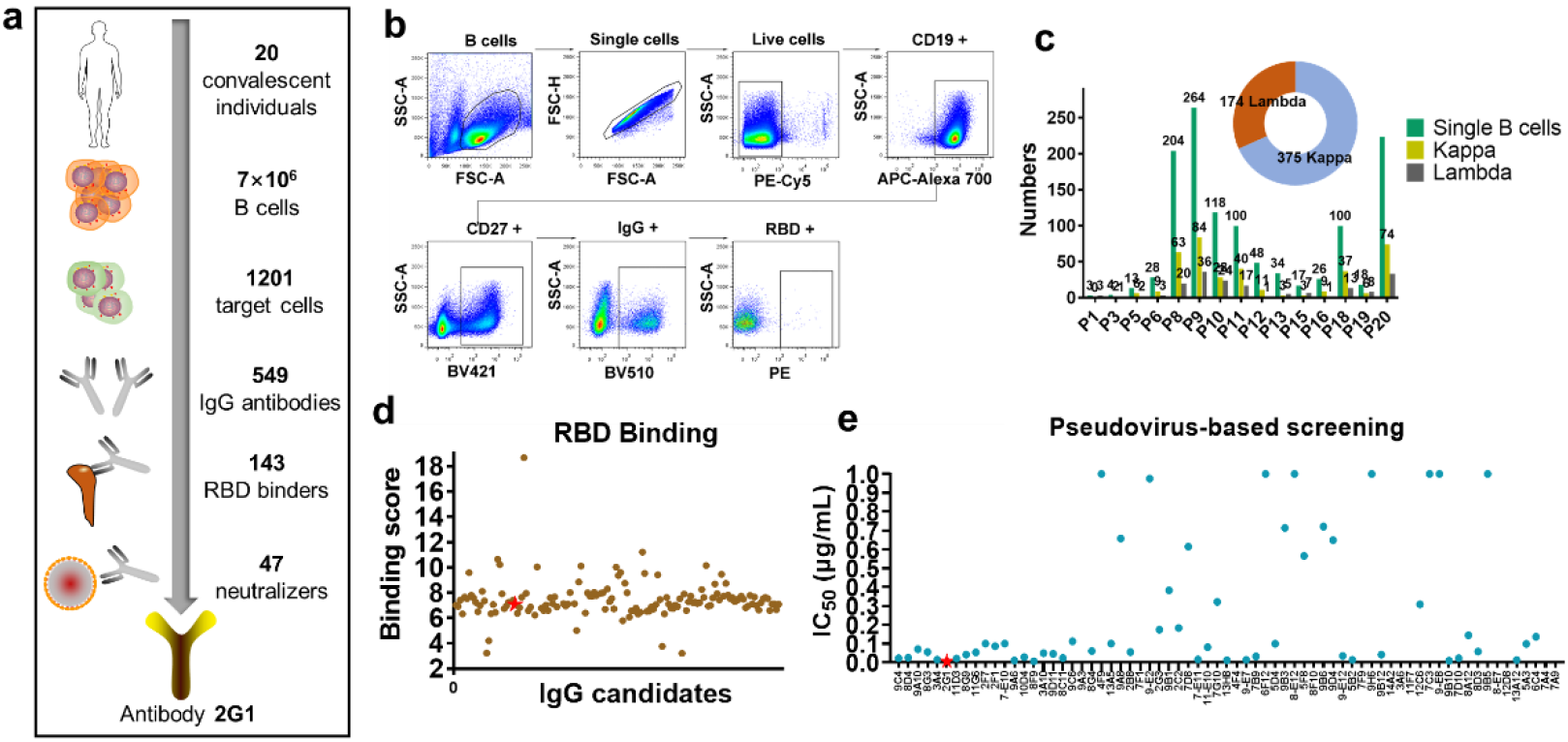
Cell isolation, antibody cloning, and candidate panning. **a,** Isolation strategy of highly potent neutralizing antibodies as depicted by a diagram. **b,** RBD-specific B cells were isolated from convalescent subjects of SARS-CoV-2 infection by fluorescence-activated cell sorting. The 7ADD^-^/CD19^+^/CD27^+^/IgG^+^/RBD^+^ gate is shown and highlighted in the boxes. **c,** Statistics of the number of paired antibodies from each subject, as well as the number of kappa and lambda subtypes. **d,** Binding scores of antibody candidates against SARS-CoV-2 RBD as measured by ELISA and scores higher than 2 are presented. 2G1 is highlighted in red. **e,** Candidate panning using a WA1/2020 pseudovirus-based screening model. Antibodies were 10-fold serially diluted from 10^1^ μg/mL to 10^-4^ μg/mL.

## Results

### Molecule discovery of 2G1

We collected blood samples from 20 convalescent individuals who were infected by SARS-CoV-2 in February 2020. Peripheral blood mononuclear cells were enriched and sorted with fluorescently labeled recombinant SARS-CoV-2 RBD (WA1/2020) protein (Fig. 1b). Over 1200 B cells were isolated and more than 500 pairs of IgG antibody genes were cloned by single-cell PCR. Of which, 375 are kappa subtype and 174 are lambda subtype (Fig. 1c). 143 RBD binders were obtained after the ELISA-based preliminary screening (Fig. 1d). In the following pseudovirus-based screening, three molecules, including 2G1, displayed ultra-potent neutralization with IC_50_ less than 0.01 μg/mL (Fig. 1e). Antibody 2G1 stood out from these candidates after further investigation despite the binding and ACE2 blocking abilities were not remarkable (Supplementary information, Fig. S1a-b). In the germline analysis of 33 candidates, 23 heavy chains were from IGHV3 and 18 light chains were from IGKV1 (Supplementary information, Fig. S2). Six heavy chains, including 2G1, were from IGHV3-53, which was reported having short complementarity-determining region and with minimal affinity but high efficacy^32^.

WA1/2020 RBD-mFc and S trimer proteins and pseudovirus were employed to further confirm the antigen-binding and neutralizing ability of 2G1. Antibody 2G1 bound to RBD-mFc and S trimer with EC_50_ of 0.016 μg/mL and 0.135 μg/mL (Fig. 2a-b) and neutralized WA1/2020 pseudovirus with IC_50_ 0.0031 μg/mL (Fig. 2c), in line with the results of previous screening. Affinity of monovalent 2G1 (Fab) to RBD was measured by surface plasmon resonance (SPR). Relatively moderate dissociation constant (K_d_) of 2G1 to WA1/2020 RBD was determined as 1.05 × 10^-3^ s^-1^. The rapid binding of 2G1 with association constant K_a_ = 2.55 × 10^6^ Ms^-1^ offered a sub-nanomolar equilibrium dissociation constant (K_D_) value of 0.41 nM (Fig. 2d). Next, the antibody 2G1 was moved to further characterization including *in vitro* and *in vivo* biological activities as well as structural and mechanism investigation.

**Fig. 2.**
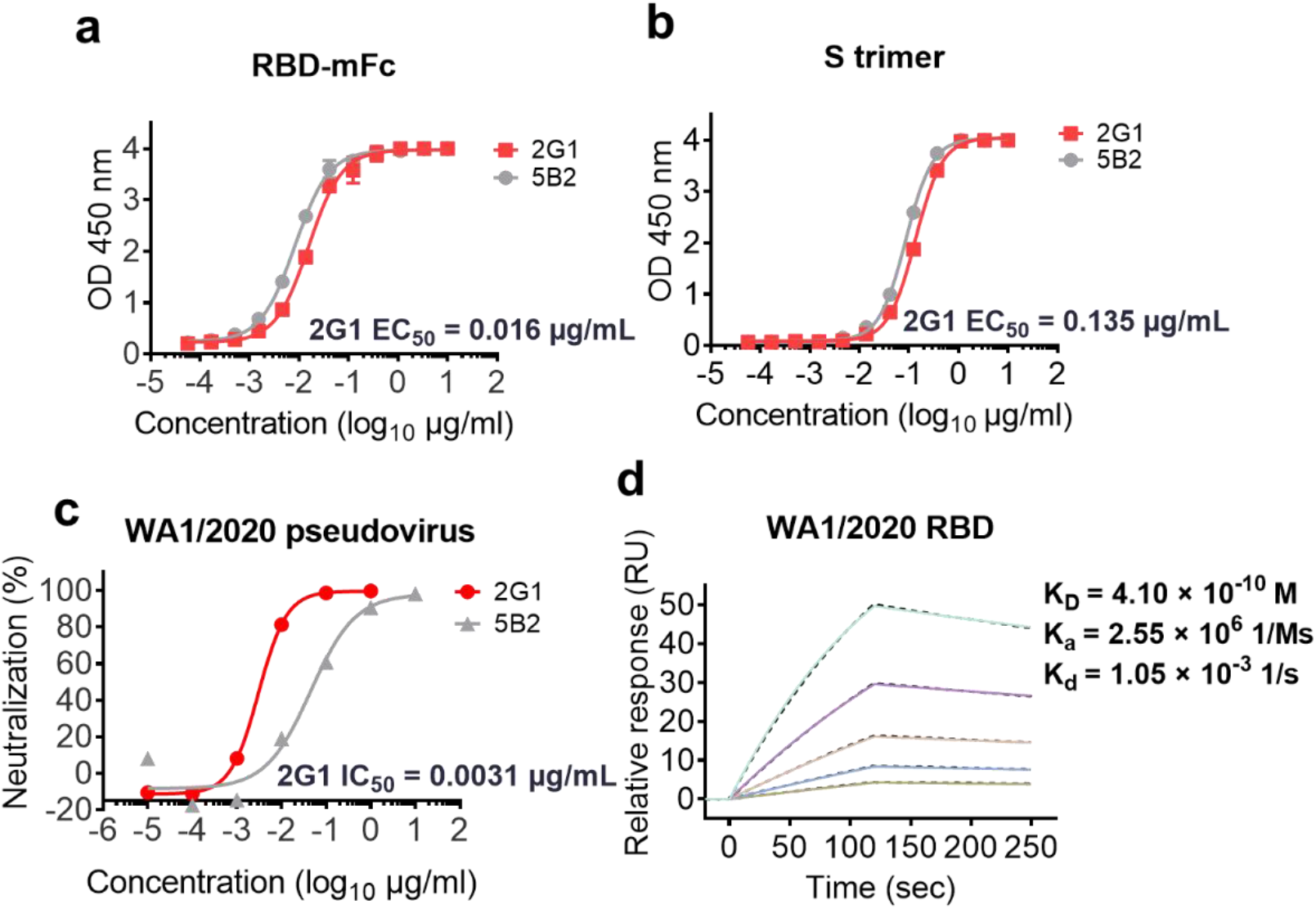
Characterization of 2G1 using WA1/2020 related S and RBD proteins and pseudovirus. **a-b,** 2G1 concentration-dependently binds to RBD-mFc (**a**) and S trimer (**b**) of SARS-CoV-2 in ELISA test. A neutralizing antibody 5B2 targeting SARS-CoV-2 RBD was used as control. Values from two replicates are shown as mean ± S.D. **c,** Serial ten-fold-diluted 2G1 was incubated with SARS-CoV-2 WA1/2020 pseudovirus and used to infect 293T-ACE2 cells. After a 48 h incubation, the infection was quantified using a fluorescence detection kit. **d,** Binding kinetics of 2G1 to SARS-CoV-2 RBD in SPR. Serial dilutions of 2G1 Fab were flowed through a chip fixed with RBD recombinant protein. The kinetics data were fitted with results from different concentrations.

### 2G1 neutralizing SARS-CoV-2 variants

With the continuing spread of mutations, combating SARS-CoV-2 variants has become a crucial task. We explored the effects of 2G1 on the mutations at several important sites such as N439K, Y453F, E484K and N501Y in terms of blocking the ACE2-RBD interaction. The IC_50_ of 2G1 blocking WA1/2020 RBD, N439K, Y453F, E484K and N501Y interacting with ACE2 were 0.1504, 0.1050, 0.2225, 0.1951 and 0.1672 μg/mL, respectively (Fig. 3a). To further study the S mutants of VOCs influence on blocking ability of 2G1, mutant trimeric S proteins of VOCs were used in ACE2 blocking experiment. The IC_50_ of 2G1 were 0.0821, 0.1066, 0.1074, 0.1047, and 0.7973 μg/mL, corresponding to WA1/2020, Alpha, Beta, Gamma, and Delta (Fig. 3b). We determined the affinities of 2G1 with various S trimers using SPR. 2G1 Fab bound to S trimers with nanomolar affinities. K_D_ of its binding to WA1/2020, Alpha, Beta, Gamma, Kappa, and Delta were 1.02, 0.86, 2.77, 2.30, 1.04, and 15.30 nM, respectively (Fig. 3c). The dissociation rate of 2G1/Delta (K_d_ = 4.27 × 10^-2^ s^-1^) was increased as compared with WA1/2020 (K_d_ = 1.05 × 10^-3^ s^-1^), which leads to the decrease in affinity.

**Fig. 3.**
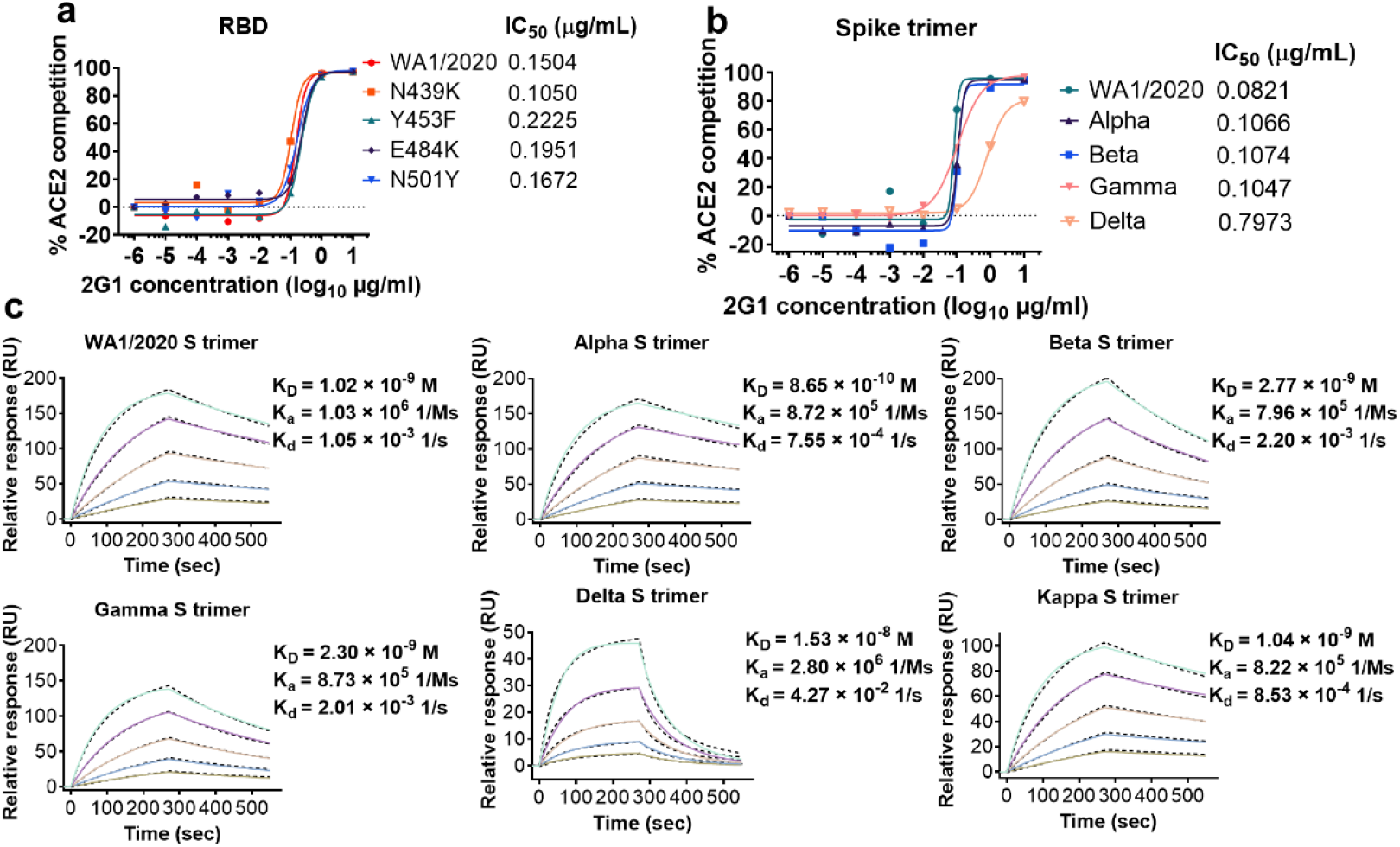
Binding and blocking characteristics of 2G1 to SARS-CoV-2 variants. **a-b,** 2G1 competitively blocked the ACE2 binding to single point mutant RBD proteins (**a**) and VOC S trimers (**b**). **c,** Affinity analysis of 2G1 bound to S trimers of SARS-CoV-2 WA1/2020, Alpha, Beta, Gamma, Kappa and Delta by SPR. Chips fixed with S trimers were loaded on a BIAcore 8K system. 2G1 Fab varied from 1.250 μg/mL to 0.039 μg/mL were injected over the chips for measuring the real-time association and dissociation parameters.

In pseudovirus neutralization assays, we found that antibody 2G1 robustly neutralized all pseudoviruses, including D614G, Alpha, Beta, Gamma, Delta, and Cluster 5 variants (Fig. 4a-g, Supplementary information, Fig. S3) with low IC_50_, especially 0.0005 μg/mL against Gamma and 0.0002 μg/mL against Cluster 5. Live SARS-CoV-2 neutralization assay results were consistent with those from experiments using pseudoviruses. Antibody 2G1 neutralized WA1/2020 live virus with IC_50_ of 0.0240 μg/mL (Fig. 4h) while it was more inclined to neutralize Alpha, Beta, and Gamma live virus, with IC_50_ decrease about 1.7-fold (0.0138 μg/mL), 5.2-fold (0.0046 μg/mL), and 3.0-fold (0.0079 μg/mL). In this assay, 2G1 had the same neutralizing activity (IC_50_ = 0.0240 μg/mL) against Delta and WA1/2020.

**Fig. 4.**
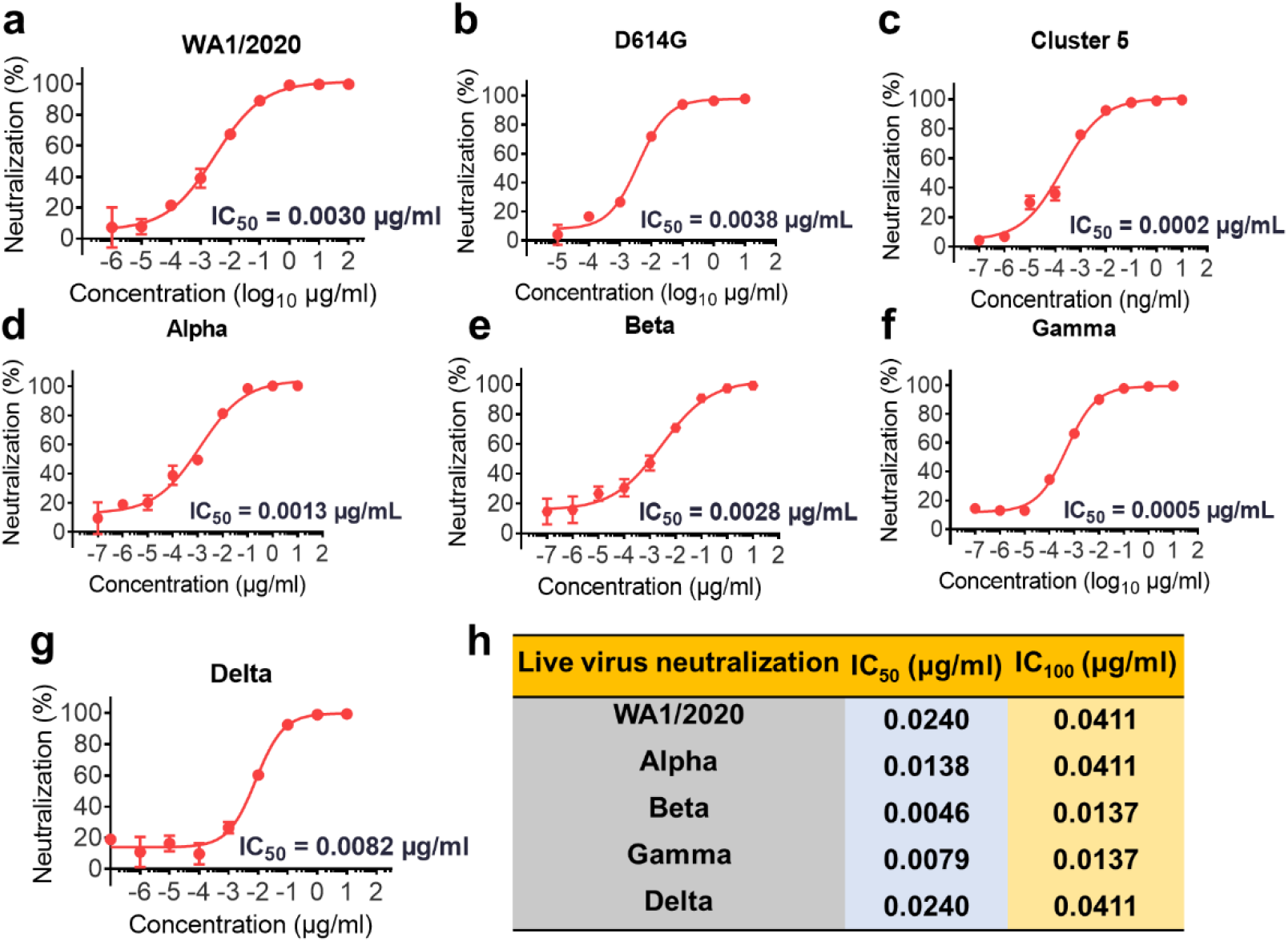
Extensive neutralization of 2G1 against SARS-CoV-2 variants. **a-g,** Neutralization of 2G1 to diverse SARS-CoV-2 pseudoviruses. Pseudoviruses with active titer higher than 1×10^7^ TU/mL were employed in this study. Concentration-dependent neutralization of 2G1 was quantified by detecting the fluorescence from the luciferase reporter. Data in duplicate are displayed as mean ± S.D. **h,** Live virus neutralization by 2G1. 100 TCID_50_ of SARS-CoV-2 (WA1/2020, Alpha, Beta, gamma and Delta) were incubated with threefold-diluted 2G1 and then added to Vero E6 cells. After a 3-day incubation, cytopathic effect (CPE) was assessed by counting the plaque formation.

### *In vivo* protection in animal models

To evaluate *in vivo* antiviral efficacy of 2G1 against SARS-CoV-2 challenge, we performed viral clearance assay employing both ACE2 transgenic mouse and rhesus macaque models. In the transgenic mouse study, animals were challenged with high copies of 100 times of half lethal dose (LD_50_) of SARS-CoV-2 WA1/2020, Beta, or Delta at day 0, followed by three different 2G1 dose treatments (20, 6.7 or 2.2 mg/kg) or vehicle injection (PBS). Four days post infection (dpi), four mice in each group including vehicle and differentially treated groups were euthanized, and lungs and brains were collected for the titration of viral load (Fig. 5a). Mice treated with vehicle developed an acute wasting syndrome and quickly met the designed endpoint at 5 dpi. In contrast, WA1/2020 and Beta virus-infected mice that received 20, 6.7 or 2.2 mg/kg treatments survived without losing any weight or revealing any obvious signs of illness throughout the study (Fig. 5b-d). Delta virus-infected mice in the 20 mg/kg group all survived throughout the trial period and had a good clinical wellbeing score. In the same study, 55.6% mice in the 6.7 mg/kg group and 10% mice in the 2.2 mg/kg group recovered back to healthy physiological condition (Fig. 5b-d) from the virus challenge. The results indicated that at the range of 6.7 - 20 mg/kg 2G1 antibody treatment was effective for animals to recover from the viral infection.

**Fig. 5.**
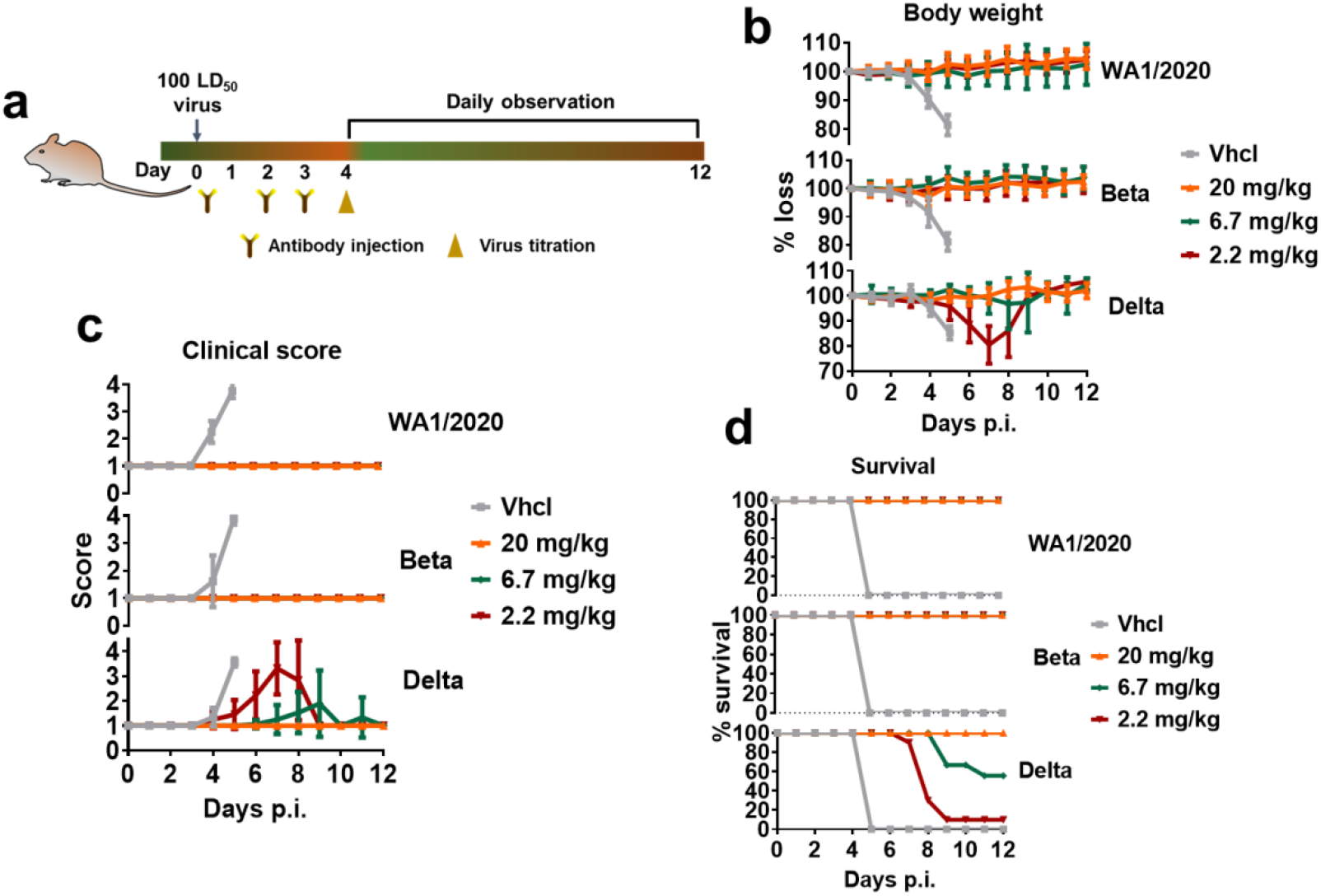
Therapeutic efficacy of 2G1 against SARS-CoV-2 variants in transgenic mice. **a,** High permissive AC70 human ACE2 transgenic mice were challenged with 100 LD_50_ of SARS-CoV-2 WA1/2020, Beta- or Delta-variants, followed by 20, 6.7, or 2.2 mg/kg of 2G1 treatment (n = 14). A 12-day clinical observation was implemented. **b,** Body weight change of mice. **c,** Clinical illness of mice was assessed based on a standardized 1 to 4 grading system that describes the clinical wellbeing of mice. **d,** Mortality of mice. Data are shown as mean ± S.D. Vhcl, vehicle control; p.i., post infection.

In the study of rhesus macaque animal model (Fig. 6a), the animals were infected with 10^5^ TCID_50_ of SARS-CoV-2 (2019-nCoV-WIV04) per animal and randomly divided into control (vehicle injection), low-dose (10 mg/kg of 2G1), and high-dose (50 mg/kg of 2G1) groups, with one male and one female in each group. Drugs were intravenously given 24 h post infection. All animals in the two therapy groups had a high viral load of 10^6^ copies/mL in the throat swab at 1 dpi. After the drug injection, the viral titer was gradually decreased. The throat virus was cleared at 3 dpi in one of the high-dose animals and at 4 dpi in the remaining treated animals (Fig. 6b). One animal in the control group had an elevated viral titer in the anal swab at 5 dpi, but no animals in the antibody treated groups showed this trend until 7 dpi (Fig. 6c). In addition, we checked the viral distribution in lung, trachea, and bronchus tissues. The virus was detectable in most areas of the lungs, in the tracheas, and bronchi of the control animals. In the group treated with high-dose of the antibody, the virus was present in right-middle, left-middle, and left-lower of the lungs, as well as left-bronchi. In the low-dose group, the virus was only found in tracheas (Fig. 6d). Results from both transgenic mouse and rhesus macaque studies showed a promising protective efficacy of 2G1, in consistent with the *in vitro* neutralization results.

**Fig. 6.**
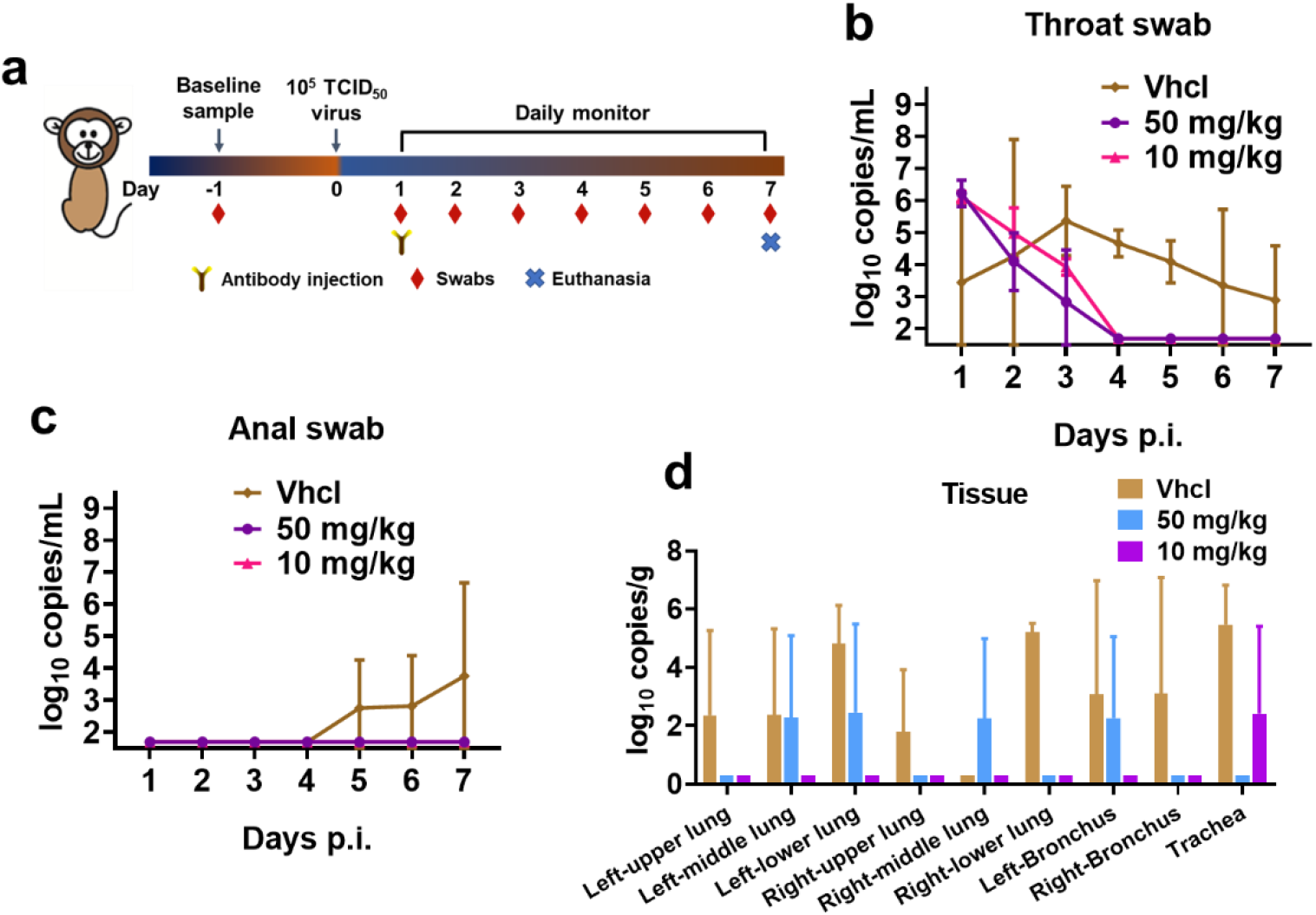
Therapeutic efficacy of 2G1 against SARS-CoV-2 variants in rhesus macaques. **a,** One male and one female rhesus macaques in each group were endotracheally challenged with 1 × 10^5^ TCID_50_ of SARS-CoV-2. 2G1 at 10 mg/kg or 50 mg/kg, or equal amount of PBS were intravenously given at 1 dpi. Throat and anal swabs were sampled daily until 7 dpi. **b,** Viral load in throat swab. **c,** Viral load in anal swab. **d,** Viral load in lungs, tracheas, and bronchi. Data with duplications are shown as mean ± S.D. p.i., post infection.

We further investigated the Fc effector function of 2G1. Results showed that 2G1 had no obvious antibody-dependent cellular cytotoxicity (ADCC) effect (not shown) but moderate antibody-dependent cellular phagocytosis (ADCP) up to 35% (Supplementary information, Fig. S4a). We hypothesize that the moderate ADCP may help the antigen presentation of SARS-CoV-2. Pharmacokinetics (PK) study revealed the half-life of 2G1 in mice was 11.1 days (Supplementary information, Fig. S4b), similar to many therapeutic antibodies. Mice treated with 15 mg/kg, 30 mg/kg, or 60 mg/kg showed no statistical changes in body weight, white blood cell count, red blood cell count, hemoglobin, and platelets (Supplementary information, Fig. S4c-g). Mice received 2G1 treatment had no evident pathological changes in hearts, livers, spleens, lungs and kidneys (Supplementary information, Fig. S5). Currently, Investigational New Drug-directed systematic safety assessment is ongoing to support the pre-clinical safety of using 2G1 in human clinical trials. Toxicology study in non-human primate showed that 2G1 was well tolerated at the maximum experimented dosage of 200 mg/kg.

### Cryo-EM structure of the complex between 2G1 and SARS-CoV-2 S protein

To investigate the binding mode of antibody 2G1 on S trimer, we solved the cryo-electron microscopy (cryo-EM) structure of 2G1 in complex with S trimer at 2.7 Å resolution (Fig. 7a, Supplementary information, Fig. S6–7). Yet, the cryo-EM map density on the interface between RBD and 2G1 were smeared. So, we performed local refinement and improved the antibody-antigen interface resolution to 3.2Å, enabling reliable analysis of the interactions between the RBD and 2G1 (Fig. 7b). In the S/2G1 complex, three solved Fabs bound to trimeric S with all RBDs in the “down” position and the S protein in a locked conformation^33,34^ (Fig. 7a). There is an additional density in RBD domain of the structure, which was reported as free fatty acid linoleic acid (LA) in a locked conformation^33^.

**Fig. 7.**
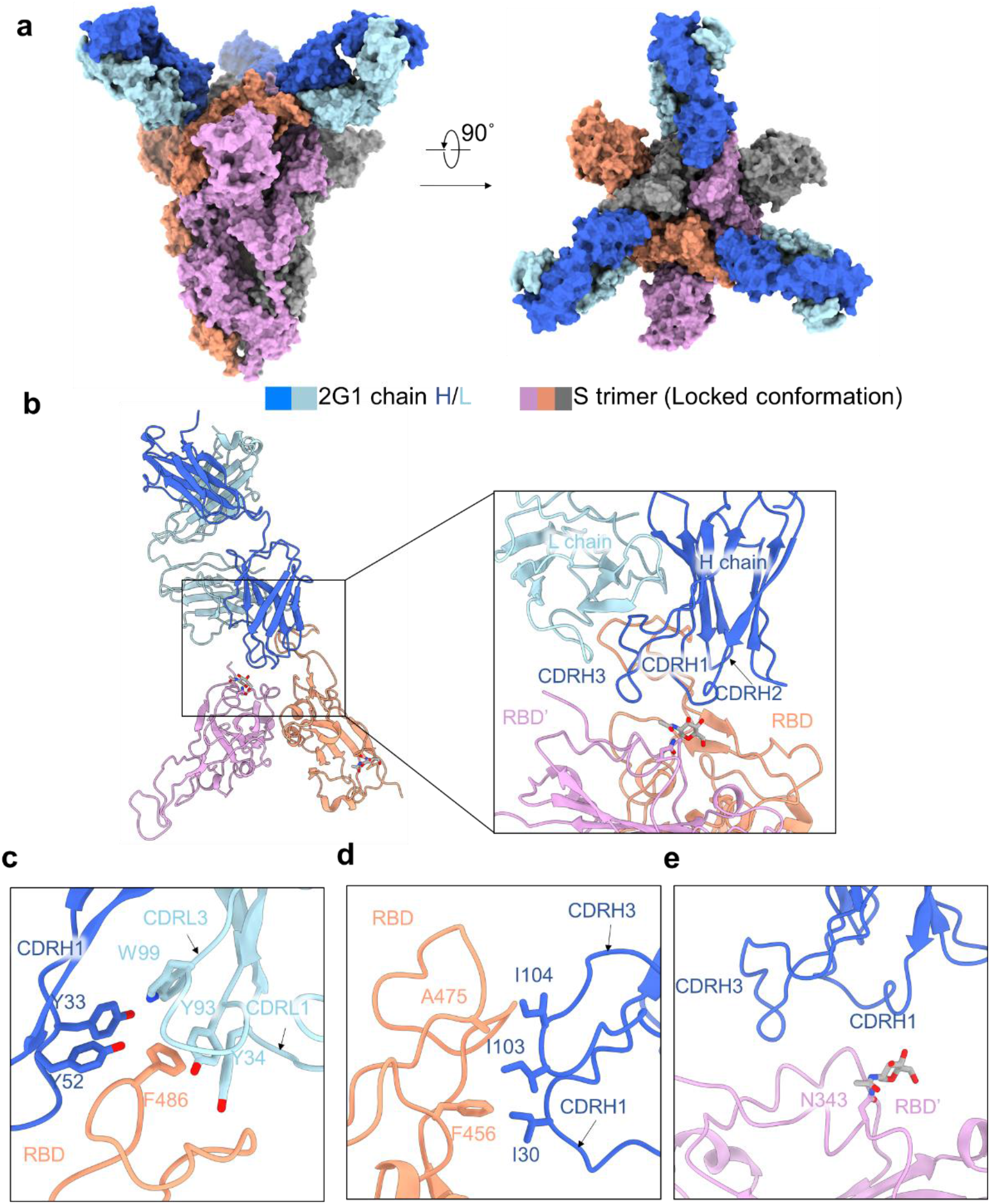
Cryo-EM structure of 2G1 and the complex with WA1/2020 S protein. **a,** The domain-colored cryo-EM map of SARS-CoV-2 S ectodomain trimer and 2G1 Fab fragments complex is shown, viewed along two perpendicular orientations. The heavy and light chains of 2G1 are colored blue and cyan, respectively. **b,** The three protomer of trimeric S protein are colored grey, orange and pink. **c-e,** The binding interface between 2G1 and RBD and adjacent RBD’. RBD and 2G1 interact each other mainly through hydrophobic interactions (**c** and **d**). 2G1 heavy chain (CDRH3 and CDRH1) lie above the adjacent RBD’ (**e**).

For detailed analysis of the interface, antibody 2G1 binds to tip area of RBD of S trimer, overlapping with the ACE2 binding site on RBD and offset from the major mutational hotspots in VOCs. The heavy chain of 2G1 interacts with RBD mainly through three complementarity-determining regions, named CDRH1 (residues 30 to 35), CDRH2 (residues 50 to 65), and CDRH3 (residues 98 to 111). The light chain of 2G1 participates interaction mainly through two CDRs, CDRL1 (residues 23 to 36) and CDRL3 (residues 91 to 100) (Fig. 7b-e). The interface between RBD and 2G1 is stabilized by an extensive hydrophobic interaction network. Phe486 on the RBD top loop interacts with Tyr33, Tyr52 on heavy chain and Tyr34, Tyr93, Trp99 on light chain through hydrophobic and/or π-π interactions simultaneously (Fig. 7c). CDRH1 and CDRH3 of the 2G1 heavy chain were positioned above the LA binding pocket in the adjacent RBD’ (Fig. 7b and 7e). We further compared 2G1 with three antibodies (S2E12, B1-182.1 and REGN10933), which have similar patterns of epitope (Fig. 8a-c). Structural comparison reveals that the epitope for 2G1 partially overlaps with these three antibodies (S2E12, B1-182.1 and REGN10933), but they have different binding directions (Fig. 8b). Besides, 2G1 has a relative narrow binding epitope which may result less probability of losing neutralizing activity due to viral mutagenesis (Fig. 8c).

**Fig. 8.**
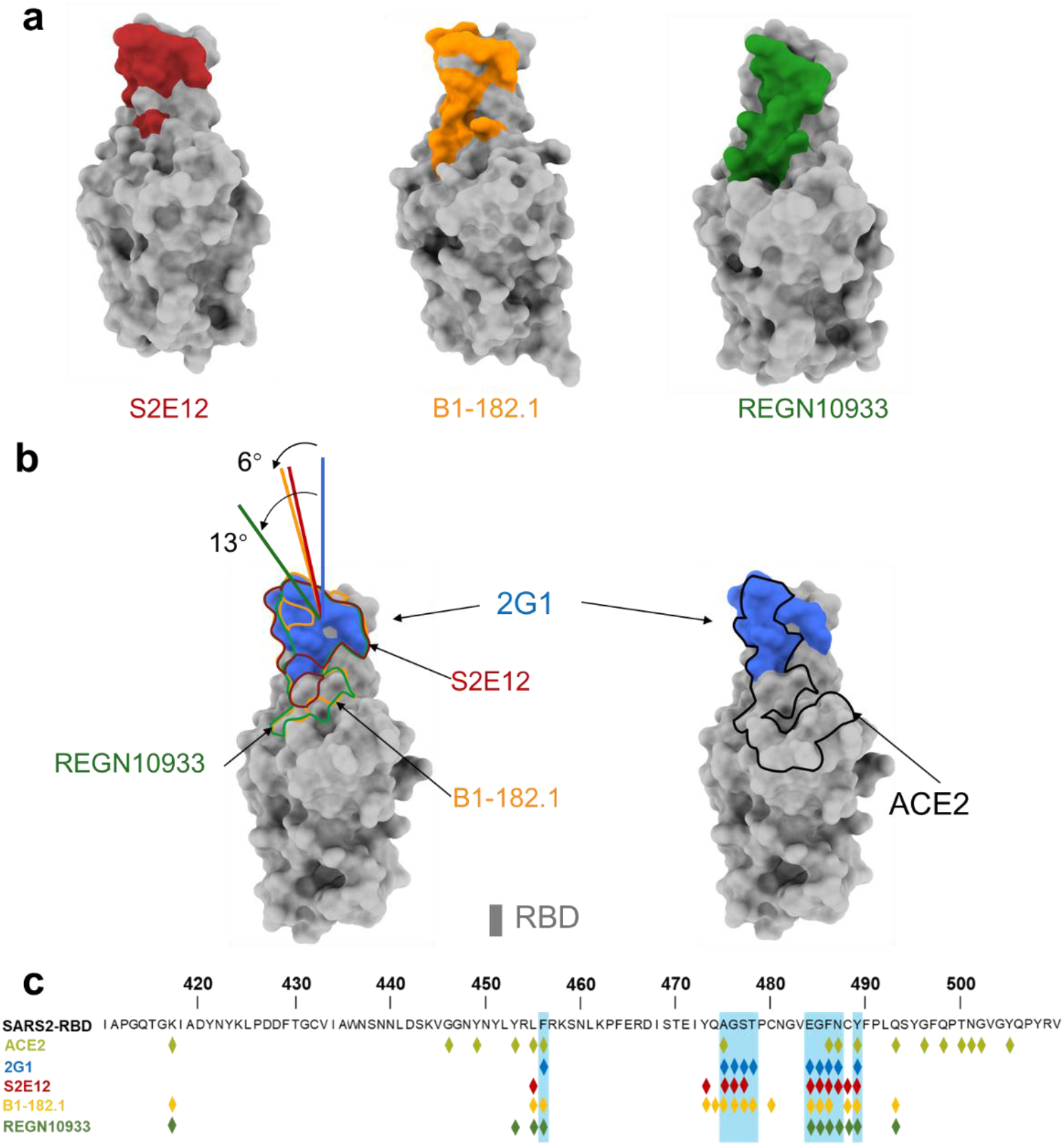
Analysis of different binding modes of 2G1, S2E12, B1-182.1 and REGN10933. **a,** The epitope surfaces of S2E12, B1-182.1 and REGN10933 on S protein are in red, orange and green, respectively. **b,** Comparison of binding modes of 2G1, S2E12, B1-182.1 and REGN10933. The epitope surface of 2G1 is in blue. The borderlines of ACE2-binding site, S2E12, B1-182.1 and REGN10933 are shown in black, red, orange and green respectively. The connecting lines between the center of 2G1 Fab and RBD is taken as the principal axis, and axis of Fab S2E12, B1-182.1 are rotated 6° and REGN10933 is rotated 13° approximately. **c,** Mapping of S2E12, B1-182.1 and REGN10933 epitopes on RBD.

### Potential escape risk evaluation

To address the potential virus escape issue, we collected the high-frequency mutation sites near the 2G1 binding epitope from GISAID database as of August 2021 (Fig. 9a), and constructed a series of S protein sequences containing these mutations. The change in binding ability of 2G1 was reflected by the normalized mean fluorescent intensity (MFI) relative to the wild-type S protein in flow cytometry. Mutants 484K, 477N/484Q/490S, and 477R/478K/484K distinctly reduced 2G1 binding (Fig. 9b). Mutants 477N/490S, 477R/490S, 478K/484Q, and 484K/490S remarkably enhanced 2G1 binding (Fig. 9b). The 484K substitution is featured in variants Beta and Gamma. Although 484K alone leads to a decreased binding ability of 2G1, trimeric S harbor all mutation sites only slightly influenced the affinity of 2G1 (Fig. 3c). The 484K substitution leads to the loss of salt bridge between Glu484 and ACE2 Lys31, resulting in the reduced affinity of ACE2^35^. It may be one of the reasons why the activity of 2G1 even slightly improved in neutralizing Beta and Gamma mutants. Another substitution in residue 484 with Gln (484Q) only slightly weakened the binding of 2G1 (Fig. 9b). SARS-CoV-2 Delta variant possesses the T478K substitution, which is a contact residue with 2G1. The single point mutation with T478K has mildly decreased the 2G1 binding (Fig. 9b), which is consistent with the SPR data.

**Fig. 9.**
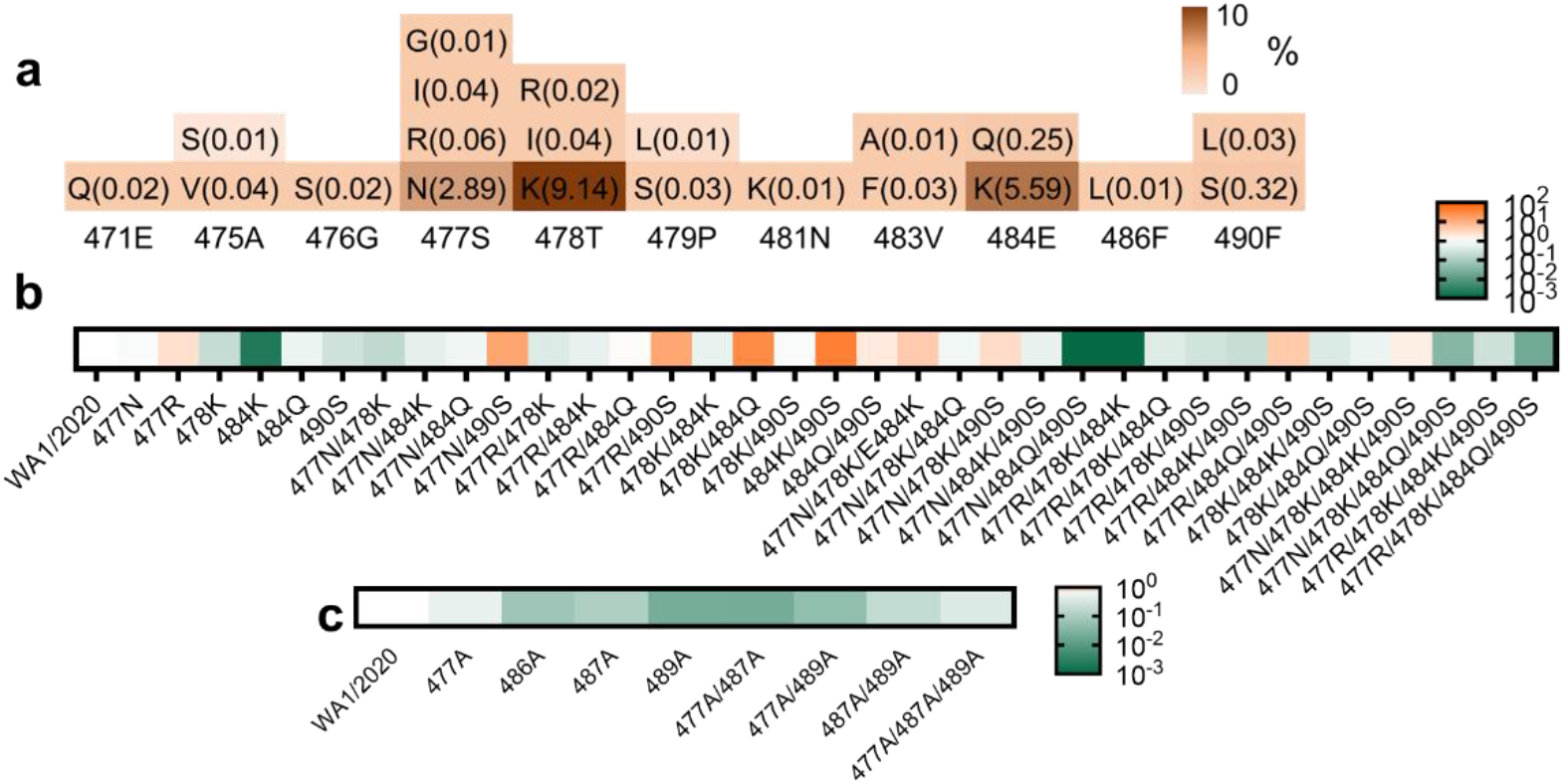
Identification of critical binding residues for 2G1. **a,** Statistics of mutation proportion in RBD residue 471Glu - 490Phe where key for 2G1 epitope from GISAID database as of August 2021. **b,** Identification of critical binding residues for 2G1. Spike genes with high frequency mutation sites between 471Glu and 490Phe (>0.05%) were cloned and transiently expressed on the surface of 293T cells. The binding ability of 2G1 to these mutant S proteins was measured by flow cytometry. The fold change of binding ability was normalized by comparing to WA1/2020 S protein. **c,** Mutations in the key interaction sites of 2G1 that affects the binding ability of 2G1 to varying degrees.

We also directly mutated the key interacting residues between RBD and 2G1 by alanine substitution, though they are not high-frequency mutation sites. Only moderate decline in 2G1 interaction was found in several mutations, including 486A, 489A, 477A/487A, and 477A/489A (Fig. 9c). These results suggest that 2G1 could potentially be effective against future SARS-CoV-2 variants.

## Discussion

SARS-CoV-2 has no sign of stopping its transmission since the outbreak, and the emergence of variants with increased transmissibility and capability of surveillance escape has assisted its continued existence. Recently, the variant Delta has become an intensively concerned strain due to its unparalleled transmissibility, which is embodied in the 1000 times higher viral load than the ancestral strain of SARS-CoV-2^6,36^. The high-frequency mutation nature of SARS-CoV-2 necessitates the development of therapies with breadth^37,38^. We screened antibodies with broad spectrum of neutralizing effects from convalescent subjects. One of which, 2G1, showed excellent and extensive neutralization to both ancestral SARS-CoV-2 WA1/2020 and VOCs at sub-nanomolar IC_50_ level. In the *in vivo* study, transgenic mice infected by the WA1/2020- and Beta-were cured by antibody 2G1 at a dose as low as 2.2 mg/kg, as well as fully protected from Delta infection in the range from 6.7 to 20 mg/kg, even when animals were challenged with 100 times LD_50_ of viral load. These results indicate that 2G1 is a potent therapeutic antibody against the broad spectrum of variants currently being concerned.

The cryo-EM structure of 2G1 in complex with the S protein revealed that 2G1 binds to the tip of S trimer through small interface but strong hydrophobic effect. The strong hydrophobic effect provides high affinity for 2G1, and the K_D_ of interaction with S trimers of SARS-CoV-2 and VOCs ranges from 0.86 nM to 15.3 nM. SARS-CoV-2 variants Beta and Gamma possess E484K and N501Y substitutions, which are adjacent to the epitope of 2G1. We correspondingly detected a slight decrease in the affinity of 2G1, from 1.02 nM for WA1/2020 to 2.77 nM for Beta and 2.30 nM for Gamma. Surprisingly, 2G1 showed no compromise in activity against Beta and Gamma in both pseudo- and live-viruses, and both *in vitro* and *in vivo*. The dose of 2.2 mg/kg of 2G1 completely cleared the viral load in Beta virus challenged transgenic mice, and the efficacy of which was as good as for WA1/2020 virus challenged mice. The IC_50_ even improved in the *in vitro* live virus test, decreased from 0.0240 μg/mL against WA1/2020 to 0.0046 μg/mL against Beta and 0.0079 μg/mL against Gamma. These results suggest that changes in affinity may not ultimately determine the therapeutic effect of neutralizing antibodies, and various other factors could be involved^35,39^. In addition, the small binding epitope reduces the probability of interference between 2G1 and other RBD antibodies so that 2G1 can cooperatively work with those antibodies to achieve a synergistic effect, for better responding to immunologic evasion of SARS-CoV-2 variants.

Furthermore, the specific 2G1 antibody epitope of RBD tip is offset from mutational hot spots and increases neutralization breadth covering new-onset VOCs. Variants Lambda comprising L452Q/F490S and Mu comprising E484K/N501Y in RBD have recently raised concerns^28,29^. Although residue 490 is nearing 2G1 epitope, our results suggested that F490S did not cause significant affinity alteration. The E484K/N501Y substitution in variant Mu is also seen in Beta and Gamma. In view of the good binding and neutralization of 2G1 against Beta and Gamma, we believe that 2G1 will likely be comparatively effective against Mu. In addition, we directly mutated the amino acid residues adjacent to the epitope on RBD by 2G1, as well as several residues that directly interact with 2G1, and found that only few mutation groups may cause a significant weakening of the 2G1 binding ability. Collectively, the model of 2G1 binding to the tip of S trimer provides a good reference for developing vaccines and optimizing a better combination therapy.

The neutralizing antibody 2G1 has been manufactured under cGMP to support the Investigational New Drug application. We would believe that antibody treatment with 2G1 will bring clinical benefit to COVID-19 patients.

## Materials and Methods

### B cells

Blood samples were obtained from patients who were recovered from COVID-19 for 10 weeks and had a negative nucleic acid test. Samples with serum antibody titer over 1 × 10^6^ were chosen for the peripheral blood mononuclear cells (PBMCs) separation using Ficoll density gradient centrifugation method. B cells were enriched applying a human B Cell Isolation Kit (Stemcell). Afterwards, B cells were then stained with APC-Alex700 labeled anti-CD19 (BD), BV421 labeled anti-CD27 (BD), BV510 labeled anti-IgG (BD), Biotin labeled RBD (Sino Biological), PE labeled streptavidin (ThermoFisher) and 7AAD (BD) Single memory B cells with potential SARS-CoV-2 antibody secretion were sorted out by gating 7AAD^-^, CD19^+^, CD27^+^, IgG^+^, and RBD^+^ using a BD Aria III cell sorter with fluorescence-activated cell sorting modules. B cells were suspended into lysis buffer and quickly frozen. B cell mRNA was subsequently converted to cDNA by SuperScript III Reverse Transcriptase (Invitrogen) and V gene were rescued by PCR. Linear Cassettes were composed of CMV promoter V_H_ or V_L_ and polyA tail, and were used for expressing a small amount of antibody for preliminary screening.

### mAb preparation

Heavy chains and light chain genes were inserted separately into pcDNA3.4 and amplified in *E. coli* DH5α. PureLink^™^ HiPure Plasmid Miniprep Kit (Invitrogen) was used for low endotoxin plasmid preparation. Monoclonal antibodies were transiently expressed by co-transfecting ExpiCHO-S cells (ThermoFisher) with heavy chain and light chain plasmids using an ExpiCHO^™^ Expression System (Gibco). Cell culture was harvested after an 8-to 14-day incubation at 37°C with humidified atmosphere of 8% CO_2_ with shaking. Full-length IgG was obtained by affinity purification utilizing a Protein A chromatography column (GE Healthcare) in AKTA avant (Cytiva). For long-term storage, antibodies were kept in a solution containing 10 mM Histidine-HCl, 9% trehalose, and 0.01% polysorbate 80.

### 293T-ACE2 cells

To obtain HEK-293T cells with stable expression of ACE2 protein, a lentiviral system bearing ACE2 (Genbank ID: BAJ21180.1) gene was constructed. In brief, HEK-293T cells (ATCC) with 70% - 80% confluence in a 10 cm dish were co-transfected with 12 μg of plasmid pHIV-puro encoding RRE and ACE2 genes, 8 μg of plasmid psPAX2 encoding gag and pol, and 4 μg of plasmid VSV-G encoding G glycoprotein of vesicular stomatitis virus(VSVG) using Lipofectamine 3000 Reagent (Invitrogen). 12 h later, the medium was changed to fresh DMEM (Gibco) supplemented with 10% FBS (Gibco) for another 48 h culturing. Medium containing virus particles was harvested and concentrated using a Lentivirus Concentration Kit (Genomeditech). The concentrated virus particles were used to infect HEK-293T cells under selection pressure of 10 μg/mL puromycin (Beyotime Biotechnology). The transfection efficiency was examined by flow cytometry using S1-mFc recombinant protein (Sino Biological) as primary antibody and FITC-AffiniPure Goat Anti-Mouse IgG (Jackson) as secondary antibody. The resulting bulk transfected population was sorted on a BD FACSJazz Cell Sorter (BD) with the BD FACS^™^ Sortware. Cells with top 1% fluorescence intensity were retained and expanded for subsequent use.

### S protein over-expression cells

The coding sequence for full-length wild-type S protein (GenBank: QHD43416.1) from Met1 to Thr1273 was inserted into plasmid pHIV-puro1.0, followed by an internal ribosome entry site (IRES) and puromycin resistance gene. The lentiviruses were generated using the HEK-293T packaging system as mentioned above. 500 μL of filtered lentivirus supernatant was added in a 24-well plate with Jurkat cells (ATCC). After cell expansion and selection with 10 μg/mL puromycin for one week, the positive S expression was confirmed by flow cytometry.

### Antigen-binding ELISA

Enzyme-linked immunosorbent assays (ELISA) were applied to study the binding ability of antibodies with SARS-CoV-2 RBDs (Sino Biological) and S trimers (AcroBiosystems). Antigens were diluted with ELISA Coating Buffer (Solarbio) to 1.0 μg/mL and immobilized onto High Binding ELISA 96-Well Plate (BEAVER) with 100 μL per well overnight at 4 °C. Plates were washed 4 times with PBST (Solarbio) and blocked with 3% skim milk for 1 h at 37°C. Then, serially diluted antibodies were added 100 μL per well and incubated at 37°C for 1h. After pipetting off the unbound antibodies, plates were washed 4 times with PBST and further incubated with 100 μL per well of goat anti-human IgG (Fc specific)-Peroxidase antibody (1: 5000 dilution, Sigma) for 1 h at 37°C. After a final 4 times washing with PBST, the binding of antibodies with SARS-CoV-2 antigens were visualized by adding 100 μL peroxidase substrate TMB Single-Component Substrate solution (Solarbio) and incubating for 15 min in dark. The reaction was terminated by adding 50μL stop buffer (Solarbio) and the plates were immediately submitted to an ELISA microplate reader (TECAN Infinite M200 Pro) to measure the optical density (OD) at 450 nm. Data were analyzed with GraphPad Prism Version 9.0.0 and EC_50_ values were determined using a four-parameter nonlinear regression.

### ACE2 competition ELISA

For experiments involving the competitive binding of antibodies to SARS-CoV-2 RBD or S trimer, recombinant hACE2-Fc protein was first biotinylated using EZ-Link Sulfo-NHS-Biotin (ThermoFisher) as the instruction described. SARS-CoV-2 RBD (Sino Biological), S trimer (AcroBiosystems), mutated RBDs (Sino Biological), and mutated S trimers (AcroBiosystems) were coated onto High Binding ELISA 96-Well Plate (BEAVER). In order to obtain an optimized hACE2-Fc concentration for this experiment, the concentration-dependent binding of biotinylated hACE2-Fc to coated SARS-CoV-2 antigens was measured by performing a conventional receptor-binding ELISA. The EC80 of biotinylated hACE2-Fc was calculated by the four-parameter nonlinear fitting. Antibodies were serially diluted in 1% BSA (Sigma) and added 50 μL into the antigen coated plates. Biotinylated hACE2-Fc at EC80 concentration was subsequently pipetted into. After incubation at 37°C for 1 h, plates were 4 times washed with PBST and incubated with 100 μL of 1: 2000 diluted Ultrasensitive Streptavidin-Peroxidase Polymer (Sigma). After further washing, 100 μL TMB was added, and followed by detection of the bound hACE2 in the microplate reader. Four-parameter nonlinear regression fitting in GraphPad Prism Version 9.0.0 was applied for result analysis.

### Surface Plasmon Resonance (SPR)

The binding affinities of antibodies to SARS-CoV-2 RBD and S trimers (wild-type/B.1.1.7/B.1.351/P.1/B.1.617.1/B.1.617.2) were tested using a BIAcore 8K system (Cytiva) together with CM5 biosensor chips (Cytiva). Antigens were diluted in pH 5.0 Acetate (Cytiva) and covalently coupled on chips using an Amine Coupling Kit (Cytiva). After reaching a 70 RU coupling level, the excess antigens were washed away and the unbound sites were blocked with ethanolamine. Antibodies were 2-fold serially diluted from 1.250 μg/mL to 0.039 μg/mL in HBS-EP buffer (Cytiva) and then injected for 120 s at 30 μL/min. After that, the binding was dissociated with HBS-EP buffer for 120 s, followed by chip regeneration with pH 1.5 Glycine (Cytiva). Parameters including K_a_, K_d_ and K_D_ values were calculated employing a monovalent analyte model with BIAevaluation software.

### Pseudovirus neutralization

ACE2-293T cells were seeded in a white 96-well plate (Corning) at a density of 1 × 10^4^ cells per well one night prior to use. Serially diluted antibodies were incubated with wild-type (Yeasen) or mutant pseudoviruses (GENEWIZ) for 0.5 h at 37°C. Human ACE2-Fc or other SARS-CoV-2 RBD specific antibodies were used as a positive control to validate data collection in different panels of screening. Medium containing equal amount of pseudoviruses but no antibodies was used as blank control. The culture medium of ACE2-293T cells was removed and then replaced by the antibody-pseudovirus mixture. All operations were conducted in the BSL-2 lab in Shanghai Jiao Tong University. After an additional 48 h incubation, the luminescence of each well was measured using a ONE-Glo^™^ Luciferase Assay System (Promega) in the Infinite M200 Pro NanoQuant (TECAN). The acquired luminescence units were normalized to those of blank control wells. Dose-dependent neutralization curves were fitted using a four-parameter nonlinear regression in GraphPad Prism Version 9.0.0.

### Plaque reduction neutralization

Plaque reduction neutralization test was performed using SARS-CoV-2 WA1/2020 (US_WA-1/2020 isolate), Alpha-(B.1.1.7/UK, Strain: SARS-CoV-2/human/USA/CA_CDC_5574/2020), Beta-(B.1.351/SA, Strain: hCoV-19/USA/MD-HP01542/2021), Gamma-(P.1/Brazil, Strain: SARS-CoV-2/human/USA/MD-MDH-0841/2021), and Delta-variants (B.1.617.2/Indian, Strain: GNL-751, a recently isolated strain from Galveston County, Texas) at Galveston National Laboratory at University of Texas Medical Branch at Galveston, Texas. Briefly, antibodies were 3-fold serially diluted in MEM medium (Gibco) from 20 μg/mL for preparing the working solution. The dilutions were mixed with equal volume of 100 TCID_50_ virus in two replicates and incubated at room temperature for 1 h. The mixture was then added into a 96-well plate covered with Vero cells. Blank controls and virus infection controls were set up simultaneously. After incubation at 37°C, 5% CO_2_ for 3 days, cytopathic effect (CPE) was observed under microscope and plaques were counted for efficacy evaluation. Wells with CPE changes are recorded as “+”, otherwise recorded as “-”. IC_50_ values were calculated according to the following equation: IC_50_ = Antilog (D - C × (50 - B) / (A - B)). Where A indicates the percentage of inhibition higher than 50%, B indicates the percentage of inhibition less than 50%, C is log_10_ (dilution factor), D is log_10_ (Sample concentration which the inhibition is less than 50%.

### ACE2 transgenic mouse protection

AC70 human ACE2 transgenic mice (Taconic Biosciences) were divided into control (100 μL PBS) and treatment (20, 6.7, or 2.2 mg/kg of 2G1, 100 μL) groups, with 14 in each group. Animal studies were carried out at Galveston National Laboratory at University of Texas Medical Branch at Galveston, Texas, an AAALAC accredited (November 24, 2020) and PHS OLAW approved (February 26, 2021) high-containment National Laboratory, based on a protocol approved by the Institutional Animal Care and Use Committee at UTMB at Galveston. Mice were challenged with 100 LD_50_ of SARS-CoV-2 (US_WA-1/2020 isolate), Beta-(B.1.351/SA, Strain: hCoV-19/USA/MD-HP01542/2021), or Delta-variants (B.1.617.2/Indian, Strain: GNL-751, a recently isolated strain from Galveston County, Texas), provided through World Reference Center for Emerging Viruses and Arboviruses (WRCEVA) were used in the study. The first dose of antibody 2G1 and PBS were given 4 h post infection; and the second and third were given 2 days and 4 days post infection. Mice were clinically observed at least once daily and scored based on a 1 to 4 grading system that describes the clinical wellbeing. In the standardized 1 to 4 grading system, score 1 is healthy; Score 2 is with ruffled fur and lethargic; Score 3 is with additional clinical sign such as hunched posture, orbital tightening, increased respiratory rate, and/or > 15% weight loss; Score 4 is showing dyspnea and/or cyanosis, reluctance to move when stimulated, or ≥ 20% weight loss that need immediate euthanasia. Four mice in each group were euthanized at 4 days post infection for assessing viral loads and histopathology of lung and brain. The remaining 10 mice were continue monitored for morbidity and mortality for up to 12 days post infection.

### Rhesus macaque protection

Rhesus macaques at six to seven years old were purchased from Hubei Tianqin Biotechnology Co., Ltd. All animal procedures and operations were approved by the ethical committee of Wuhan Institute of Virology, Chinese Academy of Sciences. SARS-CoV-2 strain 2019-nCoV-WIV04 (GISAID number: EPI_ISI_402124) was isolated from the bronchoalveolar lavage fluid of a patient who was infected COVID-19 in Wuhan in December 2019. Rhesus macaques were randomly divided into control group, low-dose (10 mg/kg of 2G1) and high-dose (50 mg/kg of 2G1) groups with one male and one female in each. Animals were endotracheally infected with 4 mL of 1 × 10^5^ TCID_50_ virus. Antibody 2G1 and PBS were intravenously given 24 h after infection. Rhesus macaques were monitored for disease-related changes during the period. Body weight and temperature were measured every day, and throat swab and anal swab samples were collected for virus titrating. Animals were euthanized at 7 dpi and tissue samples were collected for virus examining. Viral RNA was extracted using the QIAamp Viral RNA Mini Kit (Qiagen). A one-step real-time quantitative PCR was used to quantify the viral RNA according to the supplier’s instructions (HiScript^®^ II One Step qRT-PCR SYBR^®^ Green Kit, Vazyme Biotech Co., Ltd) together with primers for the RBD gene (RBD-qF1: 5’-CAATGGTTAAGGCAGG-3’; RBD-qR1: 5’-CTCAAGGTCTGGATCACG-3’).

### Antibody-Dependent Cellular Phagocytosis (ADCP)

In ADCP experiment, CD14^+^ monocytes (Allcells) were cultured and differentiated for 7 days to obtain macrophage cells. Macrophages were labeled with violet dye (ThermoFisher), and Jurkat cells with stable SARS-CoV-2 S expression were labeled with CFSE dye (ThermoFisher). 75,000 Jurkat cells were added to macrophage cells in a 96-well plate in the presence of 2G1 or the isotype control antibody. After incubating at 37°C for 30 mins, the macrophages were digested and fixed with 4% paraformaldehyde, and the proportion of double-positive cell populations was analyzed by flow cytometry.

### Pharmacokinetic study and toxicity test

For the pharmacokinetic study, BALB/c mice were tail intravenously injected with 2G1 (15, 30, or 60 mg/kg), or equivalent volume of PBS. Three males and three females were in each subset. Blood samples were collected 0.5 h, 6 h, 1 d, 2 d, 4 d, 7 d, 10 d, 15 d, 21 d, and 28 d after injection. Serum 2G1 concentration was quantified using ELISA. Briefly, Mouse Anti-human IgG Lambda (SouthBiotech) at 2 μg/mL was coated in ELISA plates. Serum samples and antibody 2G1 control were added into the plates and incubated for 1 h. After washing, a Goat Anti-human Fc HRP (Sigma) was used as secondary antibody with 1:6000 dilutions. After the chromogenic reaction by the HRP substrate (Solarbio), the plates were read at 450 nm.

Crlj:CD1(ICR) mice were randomly divided into control (PBS), 15 mg/kg, 30 mg/kg, and 60 mg/kg groups for testing the *in vivo* toxicity of 2G1, with three males and three females each group. Body weight was tracked every two days. Blood samples were collected 14 days after administration and mice were subsequently euthanized for tissue damage detection. Blood indicators including white blood cell count, red blood cell count, hemoglobin, and platelets were measured in multiple automated hematology analyzer (Sysmex XT-2000iV). Pathological changes of hearts, livers, spleens, lungs and kidneys were examined by hematoxylin-eosin (HE) staining.

### Expression and purification of S protein

The prefusion S extracellular domain (1-1208 a.a) (Genbank ID: QHD43416.1) was cloned into the pCAG vector (Invitrogen) with six proline substitutions at residues 817, 892, 899, 942, 986 and 987^39^, a “GSAS” substitution (instead of “RRAR”) at residues 682 to 685 and a C-terminal T4 fibritin trimerization motif followed by one Flag tag.

This recombinant S protein was overexpressed using the HEK 293F mammalian cells (Invitrogen) at 37°C under 5% CO_2_ in a Multitron-Pro shaker (Infors, 130 rpm). For secreted S protein production, about 1.5 mg of the plasmid was premixed with 3 mg of polyethylenimines (PEIs) (Polysciences) in 50 mL of fresh medium for 15 mins before adding to cell culture, and transiently transfected into the cells, when the cell density reached 2.0 ×10^6^ cells/mL. Cells were removed by centrifugation at 4000×g for 15 mins and cell culture supernatant was collected sixty hours after transfection. The secreted S proteins were purified by anti-FLAG M2 affinity resin (Sigma Aldrich). After loading two times, the anti-FLAG M2 resin was washed with the wash buffer containing 25 mM Tris (pH 8.0), 150 mM NaCl. The protein was eluted with the wash buffer plus 0.2 mg/mL flag peptide. The eluent was then concentrated and subjected to gel filtration chromatography (Superose 6 Increase 10/300 GL, GE Healthcare) in buffer containing 25 mM Tris (pH 8.0), 150 mM NaCl. The peak fractions were collected and concentrated to incubate with mAb. The purified S protein was mixed with the 2G1 at a molar ratio of about 1:5 for one hour, respectively. Then the mixture was subjected to gel filtration chromatography (Superose 6 Increase 10/300 GL, GE Healthcare) in buffer containing 25 mM Tris (pH 8.0), 150 mM NaCl. The peak fractions were collected for EM analysis.

### Cryo-EM sample preparation, data collection and data processing

The peak fractions of complex were concentrated to about 2.5 mg/mL and applied to the grids. Aliquots (3.3 μL) of the S/2G1 complex were placed on glow-discharged holey carbon grids (Quantifoil Au R1.2/1.3). The grids were blotted for 2.5 s or 3.0 s and flash-frozen in liquid ethane cooled by liquid nitrogen with Vitrobot (Mark IV, ThermoFisher). The prepared grids were transferred to a Titan Krios operating at 300 kV equipped with Gatan K3 detector and GIF Quantum energy filter. Movie stacks were automatically collected using AutoEMation^40^, with a slit width of 20 eV on the energy filter and a defocus range from −1.2 μm to −2.2 μm in super-resolution mode at a nominal magnification of 81,000×. Each stack was exposed for 2.56 s with an exposure time of 0.08 s per frame, resulting in a total of 32 frames per stack. The total dose rate was approximately 50 e^-^/Å^2^ for each stack. The stacks were motion corrected with MotionCor2^41^ and binned 2-fold, resulting in a pixel size of 1.087 Å/pixel. Meanwhile, dose weighting was performed^42^. The defocus values were estimated with Gctf^43^.

Particles for S in complex with 2G1 were automatically picked using Relion 3.0.6^44–47^ from manually selected micrographs. After 2D classification with Relion, good particles were selected and subject to two cycle of heterogeneous refinement without symmetry using cryoSPARC^48^.The good particles were selected and subjected to Non-uniform Refinement (beta) with C1 symmetry, resulting in the 3D reconstruction for the whole structures, which was further subject to 3D auto-refinement and post-processing with Relion. For interface between S protein of SARS-CoV-2 and 2G1, the dataset was subject to focused refinement with adapted mask on each RBD-2G1 sub-complex to improve the map quality. The dataset of similar RBD-2G1 sub-complexes were combined if possible and necessary. The re-extracted dataset was 3D classified with Relion focused on RBD-2G1 sub-complex. Then the good particles were selected and subject to focused refinement with Relion, resulting in the 3D reconstruction of better quality on RBD-2G1 sub-complex. The resolution was estimated with the gold-standard Fourier shell correlation 0.143 criterion^49^ with high-resolution noise substitution^50^. Refer to Supplementary information, Fig. S6–7 and Table S1 for details of data collection and processing.

For model building of the complex of S of SARS-CoV-2 with 2G1, the atomic model of the S in complex 4A8 (PDB ID: 7C2L) were used as templates, which were molecular dynamics flexible fitted^51^ into the whole cryo-EM map of the complex and the focused-refined cryo-EM map of the RBD-2G1 sub-complex, respectively. A Chainsaw^52^ model of the 2G1 was first obtained using the 4A8 as a template, which was further manually adjusted based on the focused-refined cryo-EM map of the RBD-2G1 sub-complex with Coot^53^. Each residue was manually checked with the chemical properties taken into consideration during model building. Several segments, whose corresponding densities were invisible, were not modeled. Structural refinement was performed in Phenix^54^ with secondary structure and geometry restraints to prevent overfitting. To monitor the potential overfitting, the model was refined against one of the two independent half maps from the gold-standard 3D refinement approach. Then, the refined model was tested against the other map. Statistics associated with data collection, 3D reconstruction and model building were summarized in Supplemental information, Supplementary information, Table S1.

### Binding to S mutants on cell surface

Plasmids encoding full length SARS-CoV-2 S (GenBank ID: QHD43416.1) with one or more mutation sites were carried into HEK-293T cells using lipofectamine 3000 (ThermoFisher) according to the manufacturer’s instruction. After 48 hours, cells were disassociated from the plates using a Cell Dissociation Buffer (ThermoFisher) followed by washing with PBS. Antibody 2G1 at 10 μg/ml was added into cells for a 30 min incubation. Subsequently, cells were washed and incubated with Alexa Fluor 647 labeled Goat anti-Human IgG (ThermoFisher) for 30 mins. After final washing, signals were acquired in flow cytometer (BD) and the binding ability to S mutants were evaluated by mean fluorescent intensity (MFI).

## Acknowledgments

Authors would like to acknowledge following organizations and individuals for their assistances in the preparation of the manuscript: Professors Chuan Qin and Jiangning Liu from Institute of Laboratory Animal Sciences, CAMS & PUMC, China for their support in initial cell based assay on neutralizing activity of the antibody generated in our lab; Dr. Liang Zhang of Jecho Biopharmaceuticals Co., Ltd. for his input on potential clinical applications of the antibodies; Professor Buyong Ma of SJTU for discussion and analysis on the structural interaction between the virus and antibody; We thank the cryo EM facility and the High-Performance Computing Center of Westlake University for providing supports. This work was funded by the National Natural Science Foundation of China (81773621, 82073751 to J.Z. 32022037, 31971123, 31800139 to Q.Z.); the National Science and Technology Major Project “Key New Drug Creation and Manufacturing Program” of China (No.2019ZX09732001-019 to J.Z.); the Key R&D Supporting Program (Special support for developing medicine for infectious diseases) from the Administration of Chinese and Singapore Tianjin Eco-city to Jecho Biopharmaceuticals Ltd. Co.; Zhejiang University special COVID-19 grant 2020XGZX099 and Shanghai Jiao Tong University “Crossing Medical and Engineering” grant 20X190020003 to JZ. This work was funded by the Leading Innovative and Entrepreneur Team Introduction Program of Hangzhou, and Special Research Program of Novel Coronavirus Pneumonia of Westlake University and Tencent Foundation. We would like to express our sincere gratitude towards the generous supports from Tencent Foundation and Westlake Education Foundation.

## Author Contributions

LH, HM designed and conducted experiments on antibody binding activities, antibody neutralizing experiments using pseudovirus system and drafted manuscript. LH, HT, HZ, LW, YK, YY, HY, HuiC, JZhang, YL conducted experiments on molecular discovery from blood sample to antibodies and characterization. MW, JL, YYue designed and executed animal study on metabolic profile and toxicology. CKT, AD, KRK, BHP designed and executed *in vitro* and *in vivo* study on virus neutralizing activity. XX provided technical instructions on antibody screening from B cells. JG provided critical discussions and manuscript editing. YX, HJ coordinated project on molecular discovery, characterization, preparation, and provided critical discussions on *in vitro* and *in vivo* animal study on virus neutralization. XZ, ZW, LY, YChen coordinated blood sample collection from convalescent individuals and facilitated B-cell screening. ZW, YH, YChang, GL, GcL, JJS, LLM, ZX conducted sample preparation, quality control, and product characterization. QZ conceived the project on structure analysis. YG designed and did the cryo-EM experiments. YZ solved the cryo-EM structures and YG and YZ analyzed the cryo-EM structures and made figures. SW, HH, AW, KY, ZS, HuaC, LiZ conducted experiments on antibody expression, analytical development, and characterization. WX, SZ, TJ, conducted *in vitro* virus neutralizing assays. YB, BZ coordinate project activities and provided critical discussion. JZhu designed the overall project, organized and coordinated activities from all participating institutes, and revised manuscript.

## Conflict of Interest

We declare that none of the authors have competing financial interests.

---

**Fig. S1.**
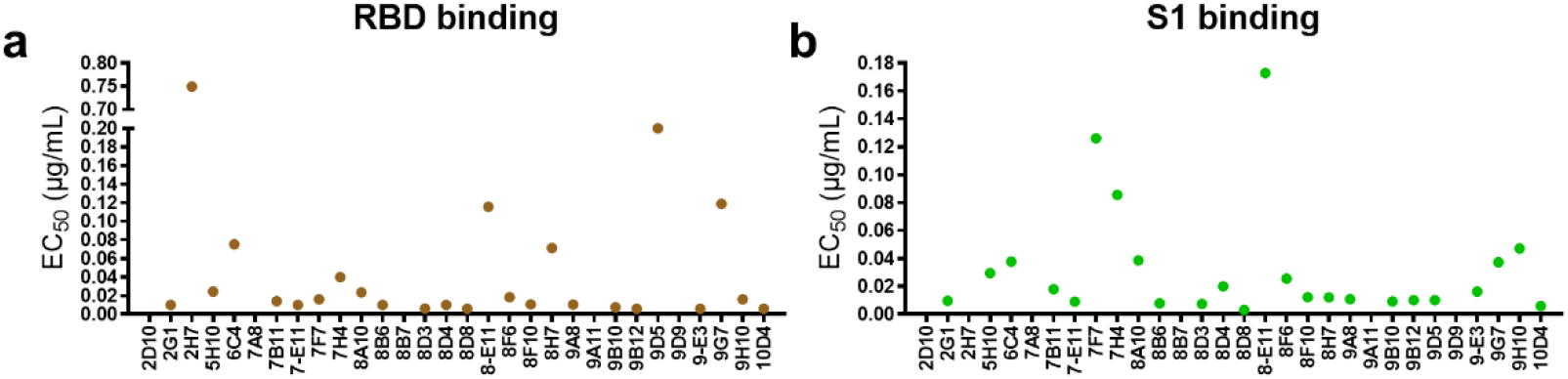
Evaluation of binding and neutralization of selected antibody candidates. **a-b,** Candidates’ EC_50_ in the concentration-dependent RBD (**a**) and S1 (**b**) binding test using ELISA. Antigens were 3-fold serially diluted from 0.300 μg/mL to 0.0012 μg/mL.

**Fig. S2.**
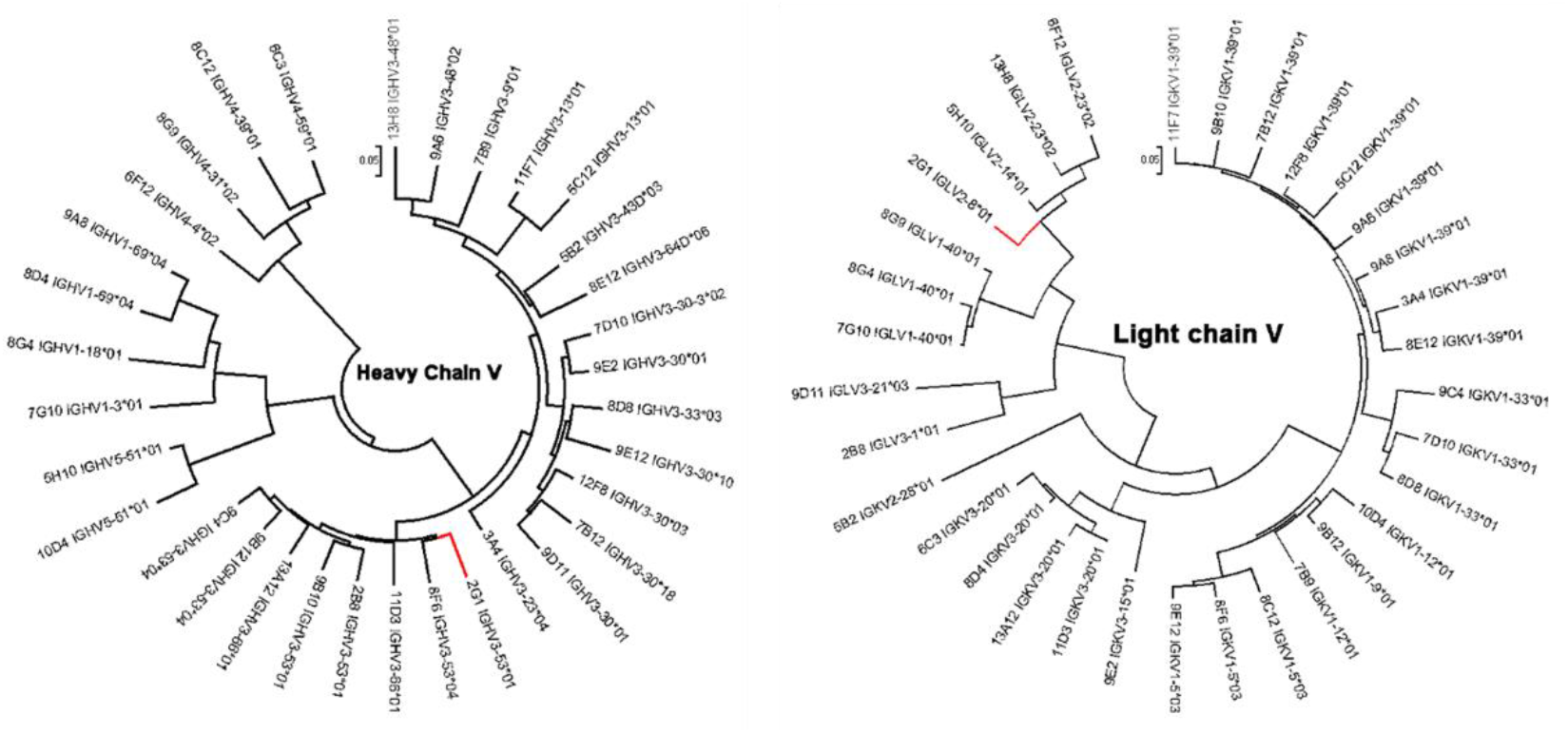
Germline identification of VH and VL. Germline gene distribution of the heavy chain and light chain of 33 candidates and their clustering analysis.

**Fig. S3.**
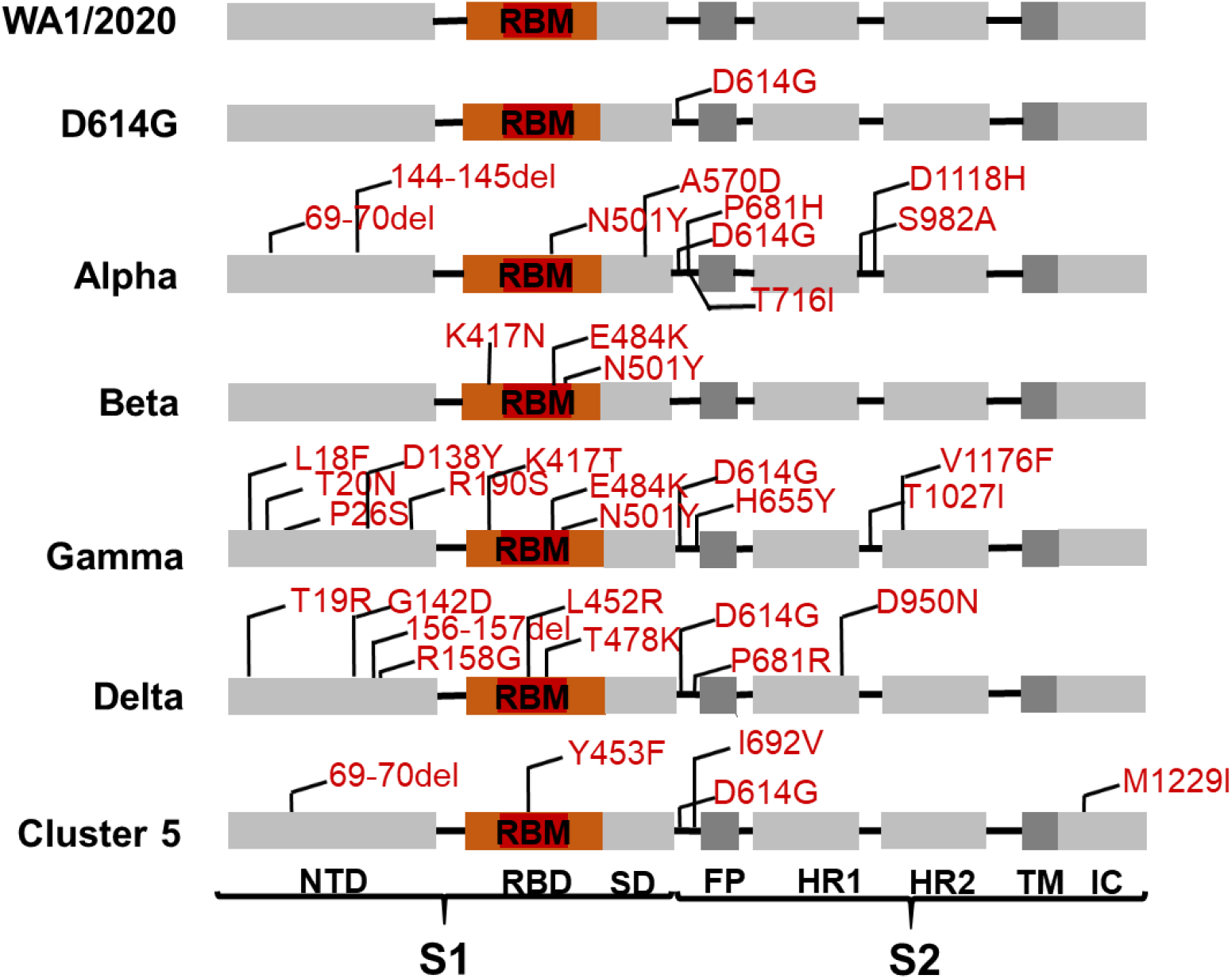
Mutational sites of pseudoviruses used in this report. The spike region of SARS-CoV-2 is displayed in different modules. The mutation sites are annotated in corresponding positions in detail. RBD is highlighted in saffron yellow and RBM is highlighted in red. NTD, N-terminal domain; RBD, receptor binding domain; RBM, receptor binding motif; SD, subdomain; FP, fusion peptide; HR1, heptad repeats 1; HR2, heptad repeats 2; TM, transmembrane region; IC, intracellular region.

**Fig. S4.**
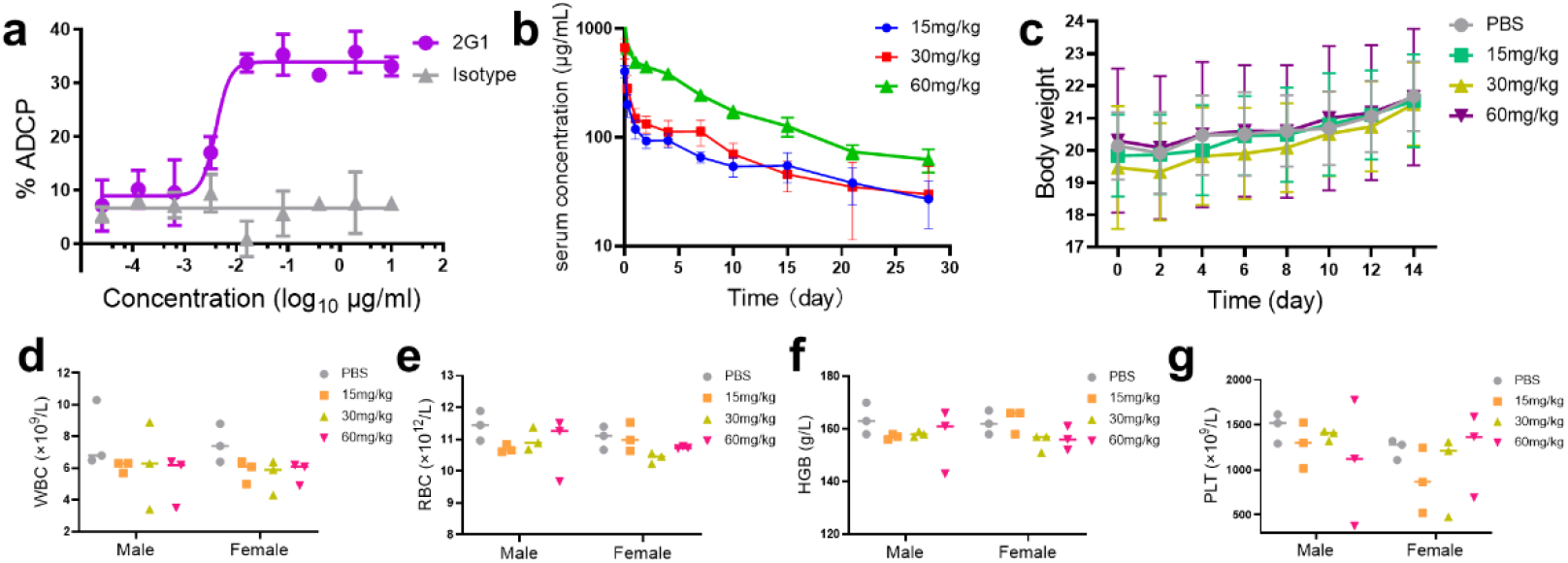
2G1 induces cellular phagocytosis but no evident adverse effects. **a,** Antibody-dependent cellular phagocytosis (ADCP) induced by 2G1. Jurkat cells with stable S expression were incubated with macrophages in the presence of different concentrations of 2G1. After incubating at 37°C for 30 mins, the proportion of Jurkat cells phagocytosed by macrophages was detected by flow cytometry. **b,** Pharmacokinetic study of 2G1. BALB/c mice were treated with different doses of 2G1, and blood samples were collected at different time points. The serum concentration of 2G1 was measured by ELISA. **c-g,** Adverse effect study of 2G1. Crlj:CD1(ICR) mice were treated with different doses of 2G1. Body weight of mice was tracked (**c**). The blood routine indexes including WBC (**d**), RBC (**e**), HGB (**f**), and PLT (**g**) were measured 14 days later. WBC, white blood cell count; RBC, red blood cell count; HGB, hemoglobin; PLT, platelets. Data are presented as mean ± S.D.

**Fig. S5.**
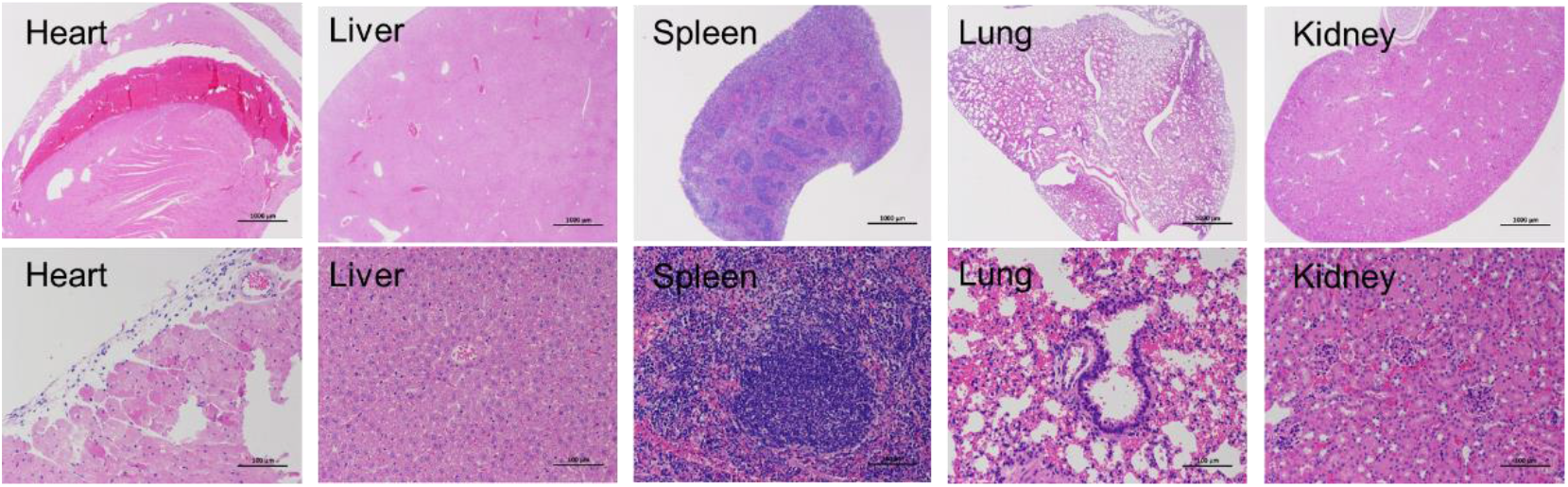
Organ toxicity study. Crlj:CD1(ICR) mice were treated with 15, 30, or 60 mg/kg of 2G1. Inflammatory damage of hearts, livers, spleens, lungs and kidneys were checked by hematoxylin-eosin (HE) staining. No apparent pathological changes were observed. Representative sections from 60 mg/kg group are displayed.

**Fig. S6.**
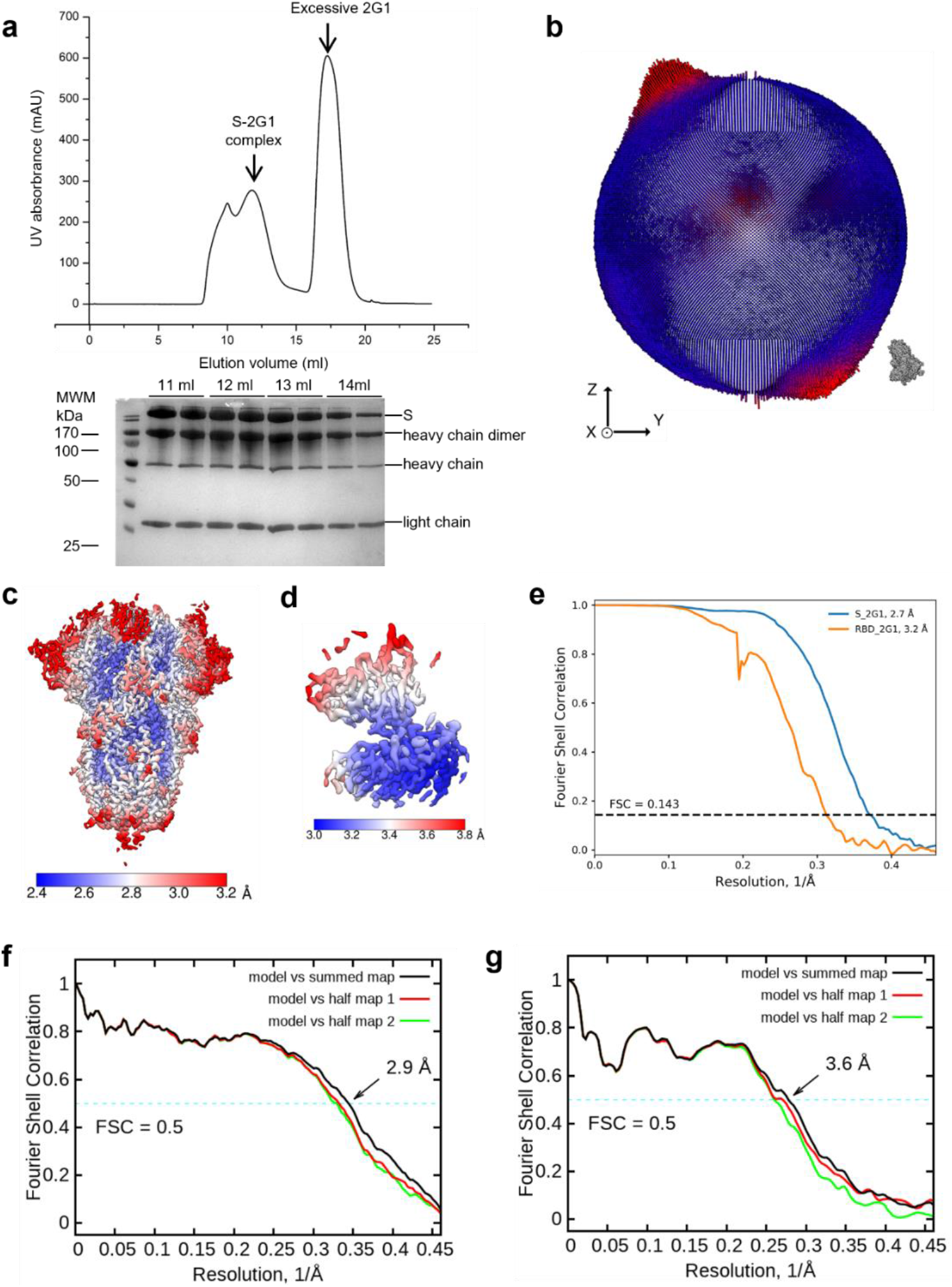
Cryo-EM analysis of SARS-CoV-2 S trimer in complex with 2G1. **a,** Representative gel filtration chromatography purification profile of the SARS-CoV-2 S extracellular domain in complex with 2G1. **b**, Euler angle distribution in the final 3D reconstruction of S bound with 2G1. **c-d,** Local resolution map for the 3D reconstruction of overall structure and RBD-2G1 sub-complex, respectively. **e,** FSC curve of the overall structure (blue) and RBD-2G1 sub-complex (orange). **f,** FSC curve of the refined model of S bound with 2G1 versus the overall structure that it is refined against (black); of the model refined against the first half map versus the same map (red); and of the model refined against the first half map versus the second half map (green). The small difference between the red and green curves indicates that the refinement of the atomic coordinates did not suffer from overfitting. **g,** FSC curve of the refined model of RBD-2G1 sub-complex, which is same to the **f**.

**Fig. S7.**
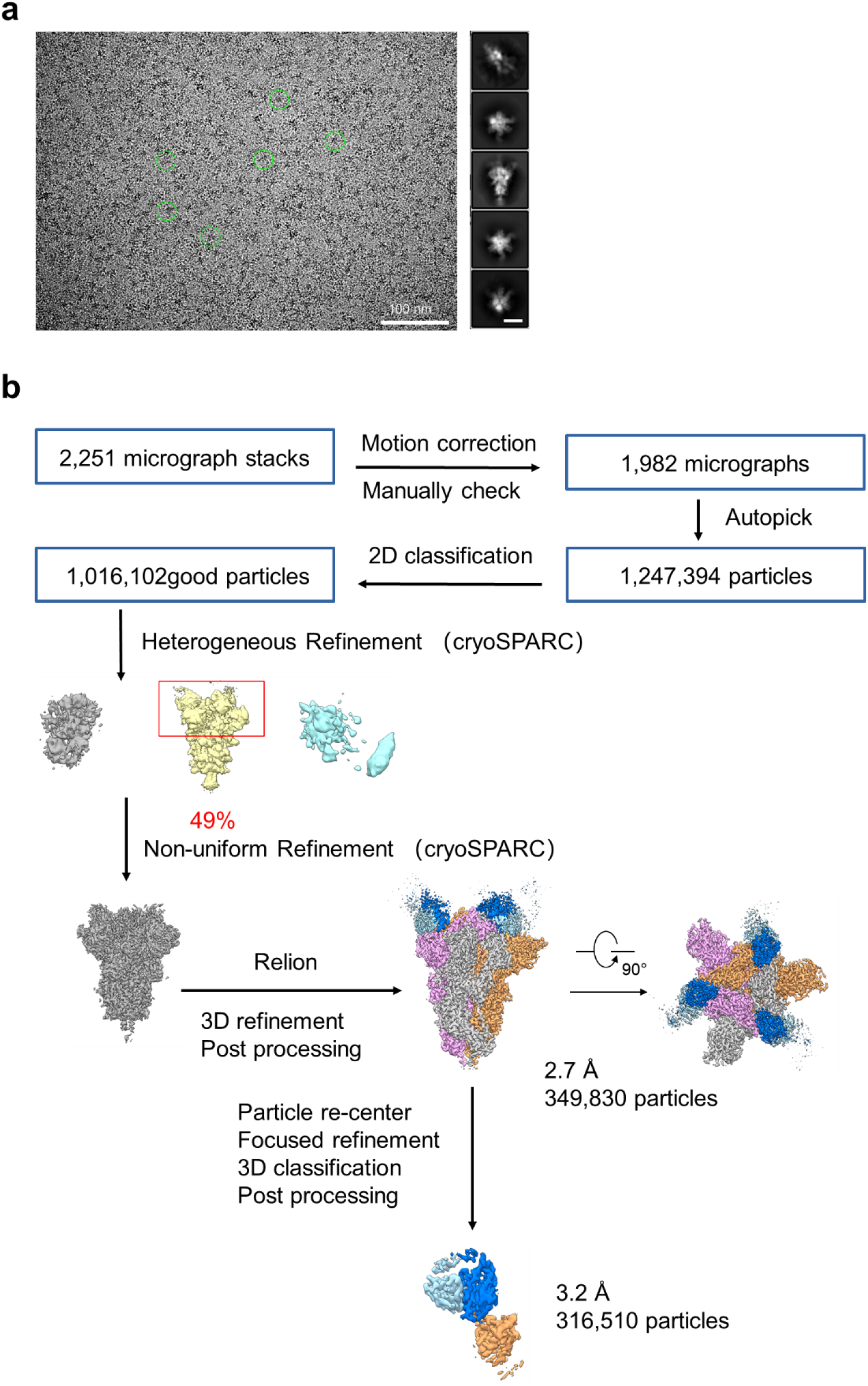
Flowchart for cryo-EM data processing of SARS-CoV-2 S trimer in complex with 2G1. **a,** Representative cryo-EM micrograph and 2D class averages of cryo-EM particle images of **SARS-CoV-2 S trimer** bound with 2G1. The scale bar in 2D class averages is 10 nm. **b,** Please refer to the ‘Data Processing’ in Methods section for details.

**Table S1.**
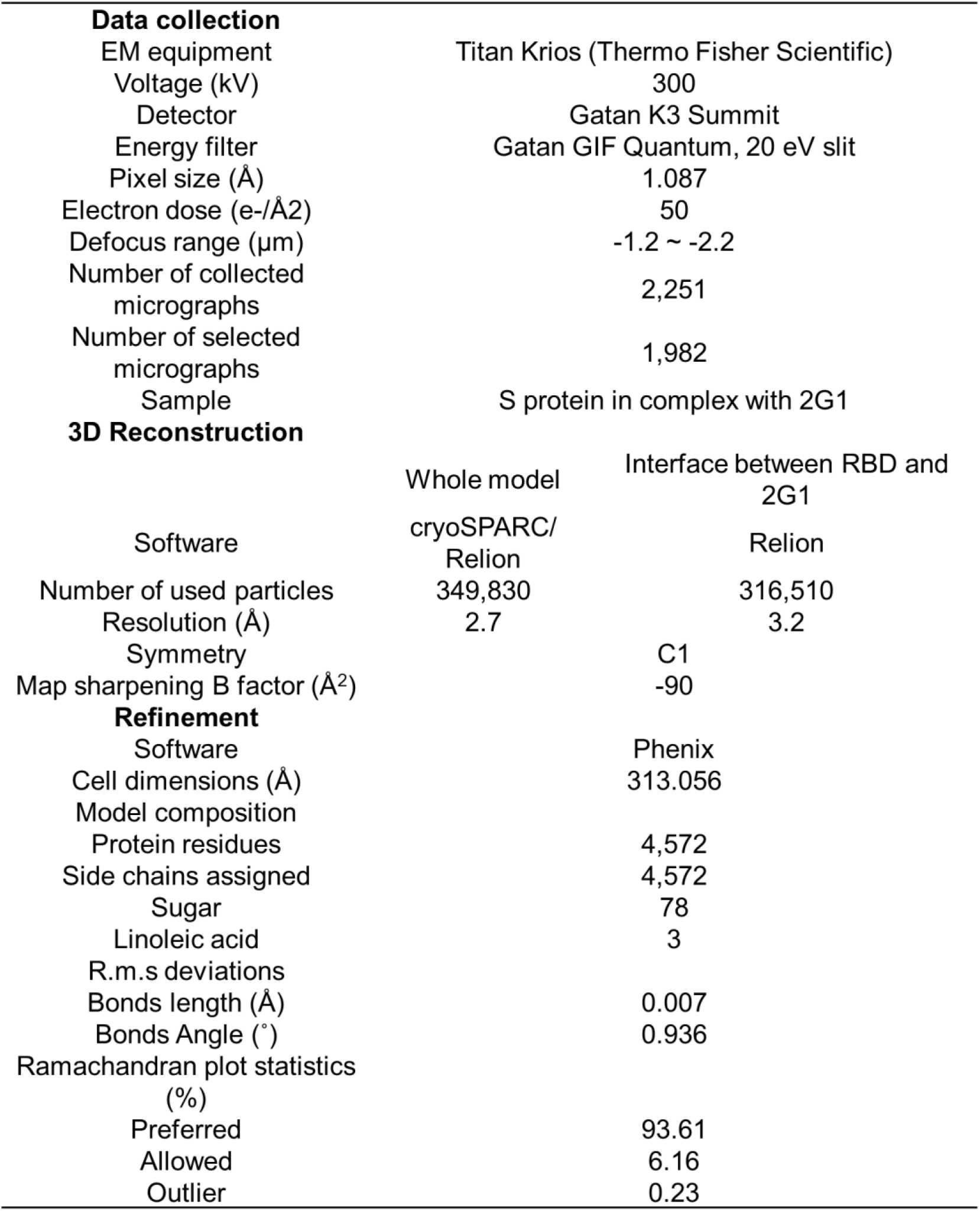
Data collection, 3D reconstruction and model statistic.

## Notes

### Competing Interest Statement

The authors have declared no competing interest.

## References

1 Volz, E. et al. Assessing transmissibility of SARS-CoV-2 lineage B.1.1.7 in England. Nature 593, 266–269 (2021).

2 Alpert, T. et al. Early introductions and transmission of SARS-CoV-2 variant B.1.1.7 in the United States. Cell 184, 2595–2604 (2021).

3 Liu, J. et al. BNT162b2-elicited neutralization of B.1.617 and other SARS-CoV-2 variants. Nature 596, 273–275 (2021).

4 Dejnirattisai, W. et al. Antibody evasion by the P.1 strain of SARS-CoV-2. Cell 184, 2939–2954.e2939 (2021).

5 Chen, R. E. et al. In vivo monoclonal antibody efficacy against SARS-CoV-2 variant strains. Nature 596, 103–108 (2021).

6 Li, B. et al. Viral infection and transmission in a large, well-traced outbreak caused by the SARS-CoV-2 Delta variant. medRxiv, doi:10.1101/2021.07.07.21260122 (2021).

7 Andreano, E. & Rappuoli, R. SARS-CoV-2 escaped natural immunity, raising questions about vaccines and therapies. Nat Med 27, 759–761 (2021).

8 Cohen, A. A. et al. Mosaic nanoparticles elicit cross-reactive immune responses to zoonotic coronaviruses in mice. Science 371, 735–741 (2021).

9 Ju, B. et al. Human neutralizing antibodies elicited by SARS-CoV-2 infection. Nature 584, 115–119 (2020).

10 Lv, Z. et al. Structural basis for neutralization of SARS-CoV-2 and SARS-CoV by a potent therapeutic antibody. Science 369, 1505–1509 (2020).

11 Chi, X. et al. A neutralizing human antibody binds to the N-terminal domain of the Spike protein of SARS-CoV-2. Science 369, 650–655 (2020).

12 Liu, Z. M. et al. Identification of SARS-CoV-2 spike mutations that attenuate monoclonal and serum antibody neutralization. Cell Host Microbe 29, 477–488 (2021).

13 Chen, R. E. et al. Resistance of SARS-CoV-2 variants to neutralization by monoclonal and serum-derived polyclonal antibodies. Nat Med 27, 717–726 (2021).

14 Garcia-Beltran, W. F. et al. Multiple SARS-CoV-2 variants escape neutralization by vaccine-induced humoral immunity. Cell 184, 2372–2383 (2021).

15 Gupta, R. K. Will SARS-CoV-2 variants of concern affect the promise of vaccines? Nat Rev Immunol 21, 340–341 (2021).

16 Kannan, S., Shaik Syed Ali, P. & Sheeza, A. Evolving biothreat of variant SARS-CoV-2 - molecular properties, virulence and epidemiology. Eur Rev Med Pharmacol Sci 25, 4405–4412 (2021).

17 Bal, A. et al. Two-step strategy for the identification of SARS-CoV-2 variant of concern 202012/01 and other variants with spike deletion H69-V70, France, August to December 2020. Euro Surveill 26, 2100008 (2021).

18 Wang, P. F. et al. Antibody resistance of SARS-CoV-2 variants B.1.351 and B.1.1.7. Nature 593, 130–135 (2021).

19 Collier, D. A. et al. Sensitivity of SARS-CoV-2 B.1.1.7 to mRNA vaccine-elicited antibodies. Nature 593, 136–141 (2021).

20 Andreano, E. et al. SARS-CoV-2 escape in vitro from a highly neutralizing COVID-19 convalescent plasma. bioRxiv, doi:10.1101/2020.12.28.424451 (2020).

21 Madhi, S. A. et al. Efficacy of the ChAdOx1 nCoV-19 Covid-19 Vaccine against the B.1.351 Variant. N Engl J Med 384, 1885–1898 (2021).

22 Zhou, D. et al. Evidence of escape of SARS-CoV-2 variant B.1.351 from natural and vaccine-induced sera. Cell 184, 2348–2361 (2021).

23 Alter, G. et al. Immunogenicity of Ad26.COV2.S vaccine against SARS-CoV-2 variants in humans. Nature 596, 268–272 (2021).

24 Hoffmann, M. et al. SARS-CoV-2 variants B.1.351 and P.1 escape from neutralizing antibodies. Cell 184, 2384–2393 (2021).

25 Planas, D. et al. Reduced sensitivity of SARS-CoV-2 variant Delta to antibody neutralization. Nature 596, 276–280 (2021).

26 Augusto, G. et al. In vitro data suggest that Indian variant B.1.617 of SARS-CoV-2 escapes neutralization by both receptor affinity and immune evasion. Allergy, doi:10.1111/all.15065 (2021).

27 Liu, C. et al. Reduced neutralization of SARS-CoV-2 B.1.617 by vaccine and convalescent serum. Cell 184, 4220–4236 (2021).

28 Padilla-Rojas, C. et al. Genomic analysis reveals a rapid spread and predominance of lambda (C.37) SARS-COV-2 lineage in Peru despite circulation of variants of concern. J Med Virol, doi:10.1002/jmv.27261 (2021).

29 Laiton-Donato, K. et al. Characterization of the emerging B.1.621 variant of interest of SARS-CoV-2. Infect Genet Evol 95, 105038 (2021).

30 Gomez, C. E., Perdiguero, B. & Esteban, M. Emerging SARS-CoV-2 Variants and Impact in Global Vaccination Programs against SARS-CoV-2/COVID-19. Vaccines-Basel 9, doi:ARTN 243 10.3390/vaccines9030243 (2021).

31 Harvey, W. T. et al. SARS-CoV-2 variants, spike mutations and immune escape. Nat Rev Microbiol 19, 409–424, doi:10.1038/s41579-021-00573-0 (2021).

32 Yuan, M. et al. Structural basis of a shared antibody response to SARS-CoV-2. Science 369, 1119–1123 (2020).

33 Toelzer, C. et al. Free fatty acid binding pocket in the locked structure of SARS-CoV-2 spike protein. Science 370, 725–730, doi:10.1126/science.abd3255 (2020).

34 Walls, A. C. et al. Structure, Function, and Antigenicity of the SARS-CoV-2 Spike Glycoprotein. Cell 181, 281–292 (2020).

35 Cai, Y. F. et al. Structural basis for enhanced infectivity and immune evasion of SARS-CoV-2 variants. Science 373, 642–648 (2021).

36 Liu, Y. & Rocklöv, J. The reproductive number of the Delta variant of SARS-CoV-2 is far higher compared to the ancestral SARS-CoV-2 virus. J Travel Med, doi:10.1093/jtm/taab124 (2021).

37 Hoffmann, M. et al. SARS-CoV-2 variant B.1.617 is resistant to bamlanivimab and evades antibodies induced by infection and vaccination. Cell Rep 36, 109415 (2021).

38 Lopez Bernal, J. et al. Effectiveness of Covid-19 Vaccines against the B.1.617.2 (Delta) Variant. N Engl J Med 385, 585–594 (2021).

39 Hsieh, C. L. et al. Structure-based design of prefusion-stabilized SARS-CoV-2 spikes. Science 369, 1501–1505 (2020).

40 Lei, J. & Frank, J. Automated acquisition of cryo-electron micrographs for single particle reconstruction on an FEI Tecnai electron microscope. J Struct Biol 150, 69–80 (2005).

41 Zheng, S. Q. et al. MotionCor2: anisotropic correction of beam-induced motion for improved cryo-electron microscopy. Nat Methods 14, 331–332 (2017).

42 Grant, T. & Grigorieff, N. Measuring the optimal exposure for single particle cryo-EM using a 2.6 angstrom reconstruction of rotavirus VP6. Elife 4, e06980 (2015).

43 Zhang, K. Gctf: Real-time CTF determination and correction. Journal of Structural Biology 193, 1–12 (2016).

44 Zivanov, J. et al. New tools for automated high-resolution cryo-EM structure determination in RELION-3. Elife 7, e42166 (2018).

45 Kimanius, D., Forsberg, B. & Lindahl, E. Accelerated Cryo-EM Structure Determination with Parallelisation using GPUs in Relion-2. Elife 5, e18722 (2016).

46 Scheres, S. H. W. RELION: Implementation of a Bayesian approach to cryo-EM structure determination. Journal of Structural Biology 180, 519–530 (2012).

47 Scheres, S. H. W. A Bayesian View on Cryo-EM Structure Determination. J Mol Biol 415, 406–418 (2012).

48 Punjani, A., Rubinstein, J. L., Fleet, D. J. & Brubaker, M. A. cryoSPARC: algorithms for rapid unsupervised cryo-EM structure determination. Nature Methods 14, 290–296 (2017).

49 Rosenthal, P. B. & Henderson, R. Optimal determination of particle orientation, absolute hand, and contrast loss in single-particle electron cryomicroscopy. J Mol Biol 333, 721–745 (2003).

50 Chen, S. X. et al. High-resolution noise substitution to measure overfitting and validate resolution in 3D structure determination by single particle electron cryomicroscopy. Ultramicroscopy 135, 24–35 (2013).

51 Trabuco, L. G., Villa, E., Mitra, K., Frank, J. & Schulten, K. Flexible fitting of atomic structures into electron microscopy maps using molecular dynamics. Structure 16, 673–683 (2008).

52 Winn, M. D. et al. Overview of the CCP4 suite and current developments. Acta Crystallogr D 67, 235–242 (2011).

53 Emsley, P., Lohkamp, B., Scott, W. G. & Cowtan, K. Features and development of Coot. Acta Crystallographica Section D-Biological Crystallography 66, 486–501 (2010).

54 Adams, P. D. et al. PHENIX: a comprehensive Python-based system for macromolecular structure solution. Acta Crystallogr D Biol Crystallogr 66, 213–221 (2010).

